# An analysis of LysM domain function in LytE when fulfilling the D,L-endopeptidase requirement for viability in *Bacillus subtilis*

**DOI:** 10.1101/2022.01.12.475998

**Authors:** Laura Uelze

## Abstract

The D,L-endopeptidase requirement states that *Bacillus subtilis* requires either the activity of the LytE or the CwlO enzyme for viability, therefore proving that these two enzymes can complement for each other despite their very different N-terminal domains. Here, we show that another D,L-endopeptidase, LytF, can also fulfill the D,L-endopeptidase requirement for viability, when expressed from the *cwlO* promoter. Both LytE and LytF contain N-terminally located LysM domains, three and five respectively. However, cells expressing another very similar D,L-endopeptidase CwlS, with four LysM domains were not viable. This led us to investigate whether a LytE protein with any one of its three LysM domains permuted can fulfill the D,L-endopeptidase requirement for viability. We found that the three LysM domains are not functionally equivalent and that the N-terminally located LysM domain plays a greater role for functioning of the LytE enzyme than the subsequent domains. Based on an investigation of orthologous enzymes in 19 *B. subtilis* species we propose an evolutionary model describing the development of the LytE-, CwlS- and LytF-type D,L-endopeptidases and their LysM domain repeats. In summary, these results show that the LytE enzyme has been optimized to fulfill the D,L-endopeptidase requirement for cell viability of *B. subtilis* with regard to the number and properties of LysM domains that mediate peptidoglycan-binding.

## INTRODUCTION

Peptidoglycan and teichoic acids are the main structural components of the *Bacillus subtilis* cell wall (Bhavsar and Brown, 2006; Barreteau *et al*., 2008; Sauvage *et al*., 2008; Vollmer *et al*., 2008; Sekiguchi and Yamamoto, 2012). Peptidoglycan is composed of glycan chains formed by polymerization of a disaccharide of *N*-acetylglucosamine (Glc*N*Ac) and *N*-acetylmuramic acid (Mur*N*Ac). Stem pentapeptides composed (proximal to distal order) of amino acids L-alanine (position 1), D-glutamate (position 2), *meso-*diaminopimelate (position 3), D-alanine (position 4) and D-alanine (position 5) extend from the lactyl group of Mur*N*Ac. The glycan strands are linked by formation of a covalent peptide bond between the carboxyl group of D-alanine (position 4) of one strand [with concomitant loss of D-alanine (position 5)] and the amino group of *meso-*diaminopimelate (position 3) of a different strand. Glycan strands can be up to 200 disaccharides in length with a stem peptide linkage index of up to ∼33%. Thus peptidoglycan forms a mesh outside the cell to which wall-teichoic acids composed of poly(ribitol- or glycerol-phosphate) are covalently attached (Weidenmaier and Peschel, 2008; Swoboda et al., 2010; Kawai et al., 2011). This rigid mesh structure confers shape on the cell and is capable of withstanding turgor pressures of up to 5 atmospheres (Sekiguchi and Yamamoto, 2012). Interference with peptidoglycan metabolism, either through inhibition of its synthesis by antibiotics (*e.g.* penicillin and vancomycin) or an imbalance between synthesis and turnover leads to aberrant cell structure, cell division and often cell death. Thus maintenance of cell wall integrity is a prerequisite for normal cell viability.

The dramatic changes that occur in cell wall structure during growth and division require exquisite control and coordination of cell wall synthesis and turnover. Autolysins play a key role in sculpting the cell wall during these processes by cleavage of glycosidic linkages in the glycan chains and peptide linkages in the stem peptides (Smith *et al*., 2000; Sekiguchi and Yamamoto, 2012). *B. subtilis* encodes a cohort of 35 enzymes with glucosaminidase, muraminidase, amidase and endopeptidase activities that confer the capability to cleave all covalent linkages of peptidoglycan thereby allowing it to be sculpted, remodeled and turned over during cell growth and division (Smith *et al*., 2000; Sekiguchi and Yamamoto, 2012). A notable feature of the autolysin cohort of *B. subtilis* is the degree of redundancy of individual enzymatic activities (Smith *et al*., 2000; Sekiguchi and Yamamoto, 2012). For example, there are seven enzymes (CwlO, CwlS, LytE, LytF, PgdS, YkfC, YqgT) with D,L-endopeptidase activity encoded in the *B. subtilis* genome, each of which cleaves the stem peptide between D-glutamate (position 2) and *meso-*diaminopimelate (position 3). However these enzymes differ in the type and/or number of domains that interact with cell wall components. Thus the LytE, CwlS and LytF D,L-endopeptidases have 3, 4 and 5 LysM domains respectively located at the amino terminus of the protein, a domain that functions to bind proteins to peptidoglycan (Mesnage *et al*., 2014). In contrast, the N-terminal domain of CwlO functions to bind the enzyme to the FtsEX ATP-type transporter that is located in the cell membrane (Yamaguchi *et al*., 2004; Dominguez-Cuevas *et al*., 2013; Meisner *et al*., 2013). Thus variance in these proteins regions allows differential localization and binding affinity of these enzymes to elements located within the cell envelope.

While none of the seven D,L-endopeptidases are essential, *lytE* and *cwlO* are synthetically lethal, signifying that the cell requires one or the other of these enzymes for viability (Bisicchia *et al*., 2007). This indicates that they perform an overlapping essential function for the cell, probably in elongation of the cell wall during growth (Bisicchia *et al*., 2007; Hashimoto *et al*. 2012). Expression of both CwlO and LytE are positively regulated by the WalRK two-component signal transduction system (TCS). In addition, LytE expression is controlled by the SigI/RsgI sigma factor, which is induced by heat and other cell wall stresses (Tseng *et al*., 2011; Liu *et al.,* 2017). Therefore the the WalRK and SigI/RsgI regulatory systems combine to fulfill the D,L-endopeptidase requirement during normal growth and stress conditions respectively.

The synthetic lethality of the CwlO and LytE enzymes is interesting on two accounts (BIsicchia *et* al., 2012). Firstly, these enzymes bind to different regions and structures within the cell envelope. The three LysM domains direct binding of LytE to peptidoglycan in the cell wall (Ishikawa *et al*., 1998; Buist *et al*., 2008; Mesnage *et al*., 2014). In contrast, the amino terminal region of CwlO binds to the FtsEX protein complex resulting in tethering of the active enzyme to the outer face of the cell membrane (Dominguez-Cuevas *et al*., 2013; Meisner *et al*., 2013). The second feature is that the high level of redundancy among the cohort of autolysins in *B. subtilis* has hampered assigning specific functions to individual enzymes. However, the requirement for D,L- endopeptidase activity evident from the synthetic lethality of CwlO and LytE permits investigation of the ability of individual enzymes with this activity to fulfill that requirement.

In this study, the ability of individual enzymes to fulfill the D,L-endopeptidase requirement for viability when individually expressed using the *cwlO* promoter is investigated. In addition, we tested the LysM domains of LytE to establish which, and how many, are needed to fulfill the D,L-endopeptidase requirement for viability. Our results show that both LytE (3 LysM domains) and LytF (5 LysM domains), but not CwlS (4 LysM domains) can fulfill the D,L-endopeptidase requirement for viability. In addition we show that the three LysM domains of LytE are not functionally equivalent, but that each alone is sufficient to fulfill the D,L-endopeptidase requirement for viability.

## MATERIAL AND METHODS

### Bioinformatic analysis

To investigate LysM domain-containing proteins, all entries containing the term ‘LysM’ were extracted from UniProt, (55,014 hits in January 2016). These data were then screened to identify *bona fide* LysM domains using the following criteria: (i) they must be from a bacterial species and be specific to an individual bacterial strain (*e.g. Bacillus subtilis* 168); (ii) putative domains must have a range of 40 – 70 amino acids (Buist *et al*., 2008; Desvaux *et al*., 2018); (iiii) each protein must have a unique amino acid. The resulting dataset contained 38,576 LysM domains located in 26,078 proteins that were encoded by 6,971 different bacterial strains in 68 phyla.

A total of 150 LysM domain-containing proteins in this dataset were identified in nineteen closely related strains of the *B. subtilis* species group (NCBI taxonomy). Cluster analysis using ClustalOmega with default settings was performed to identify the subset of proteins with a high level of identify in their catalytic domain, thereby identifying orthologues of CwlS, LytE and LytF D,L-endopeptidases orthologs in these *B. subtilis* 168 strains (Sievers *et al*., 2011; Bodenhofer *et al*., 2015). The alignment was then used for metric multidimensional scaling conducted with the bios2mds package (Pele *et al*., 2012) to identify clusters of orthologs. Similarly, the relationships among the 185 LysM domains contained in the cohort of identified orthologs was established based on an alignment of their amino acid sequence (ClustalOmega, substitution matrix = BLOSUM80), followed by construction of a phylogenetic tree using Maximum Likelihood via the phangorn package (Schliep, 2011). The best fitting amino acid substitution models were identified (LG+G+I), the different model parameters subsequently optimized and bootstrapping performed (100 replicates), before the circular phylogenetic tree was visualized with ggtree (Yu *et al*., 2016).

The phylogeny of the 19 *Bacillus subtilis* strains was inferred through alignment-free genome comparison with feature frequency profiles (FFP) (Sims *et al*., 2009). Genome sequences (excluding plasmid sequences) of all species were retrieved from GenBank. *E. coli* BL21(DE3) was included as outgroup. FPP was performed essentially as described by Wang and Ash (2015). Neighbour Joining (NJ) trees were constructed based on the Jensen-Shannon divergence matrices. It was found that varying lengths of l between 20 and 25 resulted in highly similar trees. Accounting for the higher degree of relatedness between species investigated in this study compared to Wang and Ash (2015), length was adjusted to l = 24 in the final analysis and the NJ tree visualized with ggtree (Yu *et al*., 2016). The robustness of the tree was verified with bootstrapping and bootstrapping values of 100 out of 100 were achieved for all branches.

### Strains, plasmids and growth conditions

Bacterial strains and plasmids used in this study are listed in Supporting Information Tables S2 and S3 respectively. Oligonucleotides were obtained from IDT, (Belgium) and are listed in the supporting information Table S4. *Escherichia coli* strain TG-1 (Sambrook *et al*., 1989) was used for propagating plasmids. *E. coli* strain BL21(DE3) was used for protein overexpression and purification. Strains were grown in Luria-Bertani (LB) medium and in salt-free buffered LB (Boylan *et al*., 1993), to which NaCl was added up to the desired concentration. The high phosphate defined medium (HPDM) and low phosphate defined medium (LPDM) used, are described in Botella *et al*., (2011). Antibiotics were added at concentrations described by Noone *et al*., (2014). OD_600_ was measured using a UVmini-1240 UV-VIS spectrophotometer (Shimadzu Scientific Instruments). *B. subtilis* 168 was transformed according to the procedure of Anagnostopoulos and Spizizen (1960). Growth studies were conducted with the BioTek Synergy™2 system (BioTek Instruments) at 37°C with slow shaking. Three technical repeats were performed for each growth curve.

### Strain construction

A *lytE* null mutant strain (LU1) was constructed by replacing the *lytE* gene with a spectinomycin resistance gene (*spc*) The *spc* gene was amplified from plasmid pDG1727 using primer pairs Spec_Fwd / Spec_Rev. Chromosomal regions located upstream and downstream of *lytE* were amplified by PCR using primer pairs LytE_Up_Fwd / LytE_Up_Rev_Spec and LytE_Do_Fwd_Spec / LytE_Do_Rev respectively. These DNA fragments were joined to the fragment encoding the *spc* gene by strand overlap PCR (Horton *et al*., 1989). This linear DNA fragment was then transformed into *B. subtilis* strain 168 with selection for spectinomycin resistance, thereby generating strain LU1 in which the *lytE* gene is replaced by the *spc* gene.

A second *lytE* null mutant strain (LU331) was generated in a similar manner. The strain differentiates from LC01, in that part of the *lytE* signal sequence was preserved in this mutant. The *spc* gene was amplified as described previously. Chromosomal regions located upstream and downstream of *lytE* were amplified by PCR using primer pairs LytE_Up_Fwd_2 / Signal_LytE_Rev_Spec and LytE_Do_Fwd_Spec / LytE_Do_Rev. The DNA fragments were joined to the fragment encoding the *spc* gene by strand overlap PCR and the linear DNA fragment was transformed into WT168, with selection for spectinomycin resistance, to generate strain LU331.

Strains were constructed that express the LytE protein from the *cwlO* promoter (LU17, LU21, LU25). These strains were generated through transcriptional fusion between the *cwlO* promoter and the *lytE* gene. To generate strain LU17, four PCR fragments were amplified. First, the DNA fragments located upstream and downstream of *cwlO* locus were amplified with the primer pairs CwlO_UF_Sma / CwlO_UR and QE93_Fwd_Ery / QE_94_Rev respectively. Then the *lytE* open reading frame was amplified with the primer pair LytE_DF and LytE_DR_Ery. Lastly, the erythromycin resistance gene *erm* was amplified with primer pair Mls_Fwd / Mls _Rev from plasmid pDG6464. All PCR fragments were joined together by strand overlap extension PCR and transformed into strain EB35 with selection for erythromycin resistance. The resulting strain LU17 encodes two copies of the *lytE* gene (one at the *cwlO* locus and one at the *lytE* locus). Subsequently, the *lytE* gene at the *lytE* locus was replaced with a spectinomycin resistance gene (*spc*). For this purpose, the spc gene was amplified from LU1 with the primer pair LytE_Up_Fwd / LytE_Do_Rev. The PCR fragment was then transformed into LU17 with selection for spectinomycin resistance. The resulting strain LU21 encodes one copy of the *lytE* gene at the *cwlO* locus.

Another similar strain (LU25) was generated with the additional feature that the leader region of *cwlO* was deleted, thereby stabilizing the *lytE* mRNA and yielding higher LytE protein expression (Noone *et al*., 2014). For this strain, a DNA fragment upstream of *cwlO* from strain which has a 141 bp markerless deletion of the leader region (Noone *et al*., 2014) was amplified from strain DN3000. This PCR fragment was transformed into LU21 with selection for erythromycin resistance to generate strain LU25.

In order to investigate LytE protein expression levels, the LytE protein was joined to a HA-tag which can be detected by western analysis. Strains LU45, LU47, LU61 and LU63 are derivatives of strains WT168, LS103, LU21 and LU25 in which the *lytE* gene is fused with a C-terminal HA-tag. These strains were constructed by transforming plasmid pLU2 into WT168, LS103, LU21 and LU25 with selection for kanamycin resistance. The successful integration of the plasmid into a *cwlO* mutant strain (LS103) served as proof for the functionality of the HA-tag, given that a failure would lead to a combined *cwlO* / *lytE* mutant, which is synthetically lethal (Bisicchia *et al*., 2007).

A markerless null *lytF* mutant was generated by the method of Bloor and Cranenburgh, 2006. The DNA regions upstream and downstream of *lytF* were amplified with primers LytF_Up_Fwd/ ytF_Up_Rev and LytF_Do_Fwd/LytF_Do_Rev respectively. The *cat* gene was amplified using primers Cat_Diff_Fwd/Cat_Diff_Rev and plasmid pDG1661 as template. A *dif* site for recombination was introduced into the 5’ end of each primer (Table S4). The three DNA fragments were then ligated and transformed into WT168 with selection for chloramphenicol resistance. Transformants were then grown without for 24 hours in LB medium without selection, plated onto LB agar plates and screened for loss of chloramphenicol resistance. The null mutation in *lytF* in the resultant strain LU74 (Δ*lytF*) was confirmed by PCR.

In order to generate a strain that expresses the LytF protein from the *cwlO* promoter (with *cwlO* leader – LC111, without *cwlO* leader – LC125) a similar approach as for LU21 and LU25 was followed. Downstream fragments and the erythromycin resistance cassette were amplified as described previously. The upstream fragment was amplified with the primers CwlO_UF and CwlO_UR_LytF from WT168 (containing the *cwlO* leader for LU111) or from DN3000 (no *cwlO* leader for LU125). A middle fragment (containing the *lytF* open reading frame) was amplified with the primers LytF_Fwd / LytF_Rev_Ery. The fragments were joined together by strand overlap extension PCR and the PCR product was transformed into LU74 with selection for erythromycin. The resulting strains LU111 and LU125 encode a copy of the *lytF* gene from the *cwlO* promoter (with / without *cwlO* leader) and the *lytE* gene at the *lytE* locus.

Subsequently, the *lytE* gene at the *lytE* locus was replaced with a spectinomycin resistance gene (*spc*). For this purpose, the spc gene was with the primer pair LytE_Up_Fwd / LytE_Do_Rev from chromosomal DNA of LU. The PCR fragment was then transformed into the parental strains LU111 and LU125 with selection for spectinomycin resistance. The resulting strains LU116 and LU134 encode one copy of the *lytF* gene from the *cwlO* promoter (with / without *cwlO* leader) and are knock-outs for cwlO and *lytE*.

In order to investigate LytE / LytF protein expression levels, the LytE / LytF protein were joined to a HA-tag which can be detected by western analysis. A C-terminal fusion between either the *lytE* or the *lytF* gene and a HA-tag was achieved by transforming the pLU2 plasmid into strain LU17 to generate strain LU143 and plasmid pLU6 into WT168, LU111, LU116, LU125 and LU134 to generate strains LU147, LU151, LU153, LU155 and LU157 respectively, with selection for kanamycin.

To generate strains containing a variant LytE expressed from the *cwlO* promoter in a background with endogenous *lytE* gene, the plasmids pLU16, pLU15, pLU14, pLU17, pLU18, pLU19, pLU20, pLU21, pLU22, pLU23, pLU24, pLU25, pLU26, pLU27, pLU28, pLU30, pLU31, pLU29, pLU37, pLU38 and pLU40, were transformed by double cross-over into WT168 with selection for kanamycin to generate the strains: LU239, LU207, LU212, LU216, LU220, LU223, LU227, LU232, LU235, LU261, LU265, LU269, LU272, LU276, LU280, LU286, LU288, LU312, LU314, LU315 and LU318.

To generate strains containing a variant LytE expressed from the *cwlO* promoter in a *lytE* null mutant background, the plasmids pLU16, pLU15, pLU14, pLU17, pLU18, pLU19, pLU20, pLU21, pLU22, pLU23, pLU24, pLU25, pLU26, pLU27, pLU28, pLU30, pLU31, pLU29, pLU38, and pLU40 were transformed by double cross-over into LS089 with selection for kanamycin to generate the strains: LU205, LU209, LU213, LU217, LU222, LU226, LU229, LU233, LU238, LU262, LU266, LU270, LU273, LU277, LU281, LU287, LU289, LU313, LU316 and LU320.

Strains were constructed that express a variant LytE from the *lytE* promoter (with intact *cwlO* gene). For the construction of these strains four PCR fragments were amplified. The upstream and the downstream fragments flanking the *lytE* locus were amplified with the primer pairs LytE_Up_Fwd_2 / Signal_LytE_Rev and LytE_Do_Fwd_Ery / LytE_Do_Rev from chromosomal DNA of WT168. The downstream fragment was joined to an erythromycin resistance cassette, which was amplified from pDG646 with the primers Mls_Fwd / Mls_Rev. The upstream fragment was joined to fragments of the variant *lytE* genes, which were amplified with the primers Signal_LytE_Fwd / Catalytic_LytE_Rev_Ery from the plasmids pLU40, pLU23, pLU24, pLU25, pLU20, pLU27, pLU37, pLU16, pLU17, pLU14 and pLU29. After joining of the upstream fragment (upstream homology to the *lytE* locus plus variant *lytE* gene sequence) to the downstream fragment (erythromycin resistance cassette plus downstream homology to the *lytE* locus) by strand overlap extension PCR, the linear PCR fragments were then transformed into strain LU331 with selection for erythromycin resistance, to create the strains LU335, LU337, LU339, LU341, LU372, LU374, LU398, LU399, LU401, LU403 and LU405 respectively.

In order to disrupt the *cwlO* gene in a series of strains, expressing a variant LytE from the *lytE* promoter, a fragment encompassing the *cwlO* locus was amplified with the primer pair CwlO_UF / QE94_Rev from chromosomal DNA of the *cwlO* null mutant strain EB35. The PCR fragment was then transformed by double cross-over into the strains LU335, LU337, LU339, LU341, LU372, LU374, LU399, LU401, LU403 and LU405 with selection for kanamycin resistance to obtain strains LU343, LU350, LU352, LU354, LU383, LU385, LU407, LU409, LU411 and LU413.

In order to investigate variant LytE protein expression levels, the variant LytE proteins were joined to a HA-tag which can be detected by western analysis. To enable a C-terminal fusion of the variant LytE proteins with a HA-tag, plasmid pLU46 with homology to the *lytE* catalytic domain, followed by the HA-tag and was integrated by single cross-over at the *lytE* locus in the strains LU205, LU213, LU217, LU229, LU262, LU266, LU320, LU343, LU385, LU407, LU411, LU313, LU350, LU270, LU352 and LU413 to generate the strains LU414, LU415, LU416, LU417, LU418, LU419, LU421, LU422, LU424, LU425, LU427, LU432, LU435, LU440, LU442 and LU443. Strains were selected for spectinomycin resistance.

### Plasmid construction

Plasmid pLU2, pLU6 and pLU46 enable a C-terminal transcriptional fusion between the either the *lytE* (pLU2, pLU6) or the *lytF* (pLU46) open reading frame and a HA-tag. Plasmids pLU2 and pLU6 were constructed by cloning a fragment, with homologous region to the C-terminus of the *lytE* or the *lytF* gene with the HA-tag sequence inserted before the stop codon, into the parental vector pDG780. The fragments was amplified with the following primer pair LytE_Tag_F_BamHI / LytE_Tag_R_HA_EcoRI; LytF_Tag_F_BamHI / LytF_Tag_R_HA_EcoRI and cloned into pDG780 by use of the BamHI and EcoRI restriction sites. Plasmid pLU46 is a derivative from pLU2 with a spectinomycin resistance cassette. Exchange of the antibiotic resistance cassette was achieved through amplifying the spectinomycin resistance gene from LU1 with Spec_Fwd_EcoRI and Spec_Rev_SalI and cloning it into pLU2 via the SalI and EcoRI restriction sites.

A parental plasmid (pLU10) was constructed for convenient cloning of variant *lytE* open reading frames into a cassette that would allow integration of variant *lytE* genes at the *cwlO* locus, thereby exchanging the *cwlO* against the *lytE* open reading frame. Plasmid pLU10 was generated in a two-step process: First, upstream homology (containing the *cwlO* leader and promoter and part of the *lytE* signal sequence, with a silent BamHI restriction site) was amplified with CwlO_UF_LysM and CwlO_UR_LysM and cloned into pdG780 via the NotI and BamHI restriction sites to create plasmid pLU9. Secondly, downstream homology was amplified with the primer pair CwlO_DF_LysM and CwlO_DR _LysM and cloned into the vector pLU9 via the XhoI and KpnI restriction sites resulting in plasmid pLU10.

To obtain plasmid pLU37, harbouring a variant *lytE* open reading frame without a LysM domain, a fragment spanning the signal sequence, the spacer and the sequence of the catalytic domain was amplified with the primers LytE_Spacer_Fwd_Sig / Catalytic_LytE_Rev_XmaI and cloned into pLU10 via the XmaI and BamHI sites.

Three plasmids, that harbour a variant *lytE* open reading frame with a single LysM domain each, were obtained in a similar way, following a two-step process. (i) First, the three LysM domains sequences were amplified with the primer pairs L1_Fwd_Sig_BamHl / L1_Rev_Spac, L2_Fwd_Sig_BamHl / L2_Rev_Spac and L3_Fwd_Sig_BamHl / L3_Rev_Spac. (ii) Secondly the *lytE* catalytic domain was amplified with the primer pair Catalytic_LytE_fwd_spac / Catalytic_LytE_rev_XmaI. Then the fragments generated in (i) were joined individually to the PCR fragment (ii), utilizing the spacer sequence as overlapping region. The final fragments were then cloned into pLU10 after digestion of the vector and the fragments with XmaI and BamHI to generate the plasmids pLU23 (*lytE[L1cat]*), pLU24 (*lytE[L2cat]*) and pLU25 (*lytE[L3cat]*).

Nine plasmids were generated, harbouring a variant *lytE* open reading frame with two LysM domains each. To do so, the three LysM domains of *lytE* were amplified with two primer pairs each (creating six PCR fragments in total). One primer pair generated part of the *lytE* signal sequence, followed by the LysM domain, followed by the linker sequence (L1_fwd_sig_BamHI / L1_rev_link, L2_fwd_sig_BamHI / L2_rev_link, L3_fwd_sig_BamHI / L3_rev_link). The second primer pair generated the linker sequence followed by the LysM domain, followed by parts of the spacer sequence (L1_fwd_link / L1_rev_spac, L2_fwd_link / L2_rev_spac, L3_fwd_link / L3_rev_spac). For cloning purposes, a silent BamHI restriction site was introduced into the signal sequence and a silent SpeI restriction site was integrated in the linker sequence. The fragments were then combined and joined to the *lytE* catalytic sequence as described before and cloned into the parental vector pLU10, digested with BamHI and XmaI. This process resulted in the plasmids: pLU14 (*lytE[L13cat]*), pLU15 (*lytE[L12cat]*), pLU16 (*lytE[L11cat]*), pLU17 (*lytE[L21cat]*), pLU18 (*lytE[L22cat]*), pLU19 (*lytE[L23cat]*), pLU20 (*lytE[L31cat]*), pLU21 (*lytE[L32cat]*) and pLU22 (*lytE[L33cat]*)

Plasmids for the generation of variant LytE with three LysM domains were obtained by cloning PCR fragments of single LysM domains into vectors already harbouring a *lytE* open reading frame with two LysM domains. The three individual LysM domain sequences of *lytE* were amplified with the following primer pairs: L1_fwd_ link / L1_rev_ link (L1), L2_fwd_link / L2_rev_link (L2), L3_fwd_ link / L3_rev_link (L3). The individual LysM domains could then be cloned into the parental vectors after digestion of the vectors and the fragments with SpeI. The number and orientation of integrated domains was verified by a screening PCR and sequencing. Cloning the L1 fragment in pLU20, pLU19, pLU18 and pLU14 resulted in the plasmids pLU26, pLU29, pLU31 and pLU38. Cloning the L2 fragment in pLU18 and pLU14 resulted in the plasmid pLU30 and pLU40. Cloning the L3 fragment in pLU20 and pLU19 resulted in the plasmids pLU27 and pLU28.

To create plasmids that would allow the overexpression and purification of variant LytE proteins in *E. coli*, the respective protein coding sequences (excluding the start codon and the *lytE* signal sequence) were amplified with the forward primers Exp_L1_Fwd, Exp_L3_Fwd, Exp_L1_Fwd, Exp_L1_Fwd, Exp_L3_Fwd and Exp_Spac_Fwd from the plasmids pLU40, pLU20, pLU14, pLU23, pLU25 and pLU37 to generate the plasmids pLU47, pLU49, pLU48, pLU50, pLU51 and pLU52. The same reverse primer (Exp_Cat_Rev) was used in all reactions. PCR fragments were cloned into the NdeI/XhoI sites of the vector pET21b+ thereby re-generating the start codon and generating a C-terminal fusion between the protein and a 6xHis-tag.

### RNA extraction and RT-qPCR

RNA was extracted and transcripts were quantified using RT-qPCR as previously described by Salzberg *et* al., 2013). The following primer were used to amplify the transcripts: 16s (16S_Bsu_F / 16S_Bsu_R), *cwlO* (YvcE_5′_qPCR_orf*2 / YvcE_3 ′_qPCR_orf*2), *lytE* (LytE_qPCR_F2 / LytE_qPCR_R2), *lytF* (LytF_qPCR_fwd / LytF_qPCR_rev), *mreBH* (MreBH_qPCR_F / MreBH_qPCR_R) and *sigI* (SigI_Chip_fwd / SigI_Chip_rev).

### Western analysis

Western analysis was performed as described in Noone *et al*. (2012).

### Immunofluorescence microscopy

Immunofluorescence microscopy was performed as described previously (Noone *et al*., 2012).

### Protein expression and purification

The variant LytE proteins were expressed in *E. coli* strain BL21(DE3) which harboured plasmids pLU47 to pLU53. Protein expression and purification was performed as described previously (Howell *et al*., 2006).

### Insoluble peptidoglycan pull down assay

To determine the binding capacity of variant LytE proteins to peptidoglycan an insoluble peptidoglycan pull down assay was performed as described by Wong *et al*. (2014). Purified cell wall of *B. subtilis* 168 was prepared as described in Salzberg and Helmann (2007) (the number of washes was increased to 15). Forty-four pmoles of variant LytE protein were mixed with purified cell wall of *B. subtilis* in assay buffer (50 mM Tris/HCl, pH 8, 0.05 M NaCl, 5 mM ß-mercaptoethanol, 1 % TWEEN20) in a 25 µl reaction volume. Purified cell wall was added until OD600 measurements of the mixture reached a final absorbance of 10. For a negative control, 44 pmole of the wild type LytE protein were mixed with the assay buffer, without added cell wall material. All mixtures were incubated for 10 minutes at 10°C and centrifuged for 10 minutes at 10500 g. Both, the supernatant and the pellet were collected. The pellet was resuspended in 15 µl assay buffer.

The level of LytE protein that bound to the cell wall, and that which remained in the supernated was established on a 12.5 % SDS / PAGE gel. The proteins were visualized by coomassie blue staining and the image was quantified with the ImageJ software (Schneider *et al*., 2012). To test the influence of salt on the binding of variant LytE proteins to cell walls, NaCl were added to the assay buffer at final concentrations of 0.05 M, 0.1 M, 0.2 M, 0.3 M, 0.4 M, 1 M. Two replicates of the pull down assay at 1 M NaCl were conducted to ensure the reproducibility of the assay. A negative control was included in all experiments.

## RESULTS

### Bioinformatic analysis of the LysM domains

There are seven D,L-endopeptidases encoded in the *B. subtilis* genome, three of which contain LysM domains (*cwlS*, *lytE*, *lytF*). These three enzymes are structurally very similarly and their C-terminally located catalytic domains exhibit a high sequence similarity. However, they distinguished by the number of LysM domains with LytE, CwlS and LytF having three, four and five N-terminally located domains respectively. Although the high sequence similarity of their catalytic domains indicates a close evolutionary relationship, the manner in which these three enzymes evolved to have different numbers of LysM domains is unknown. Therefore, we sought to establish the evolutionary relationship between the CwlS, LytE and LytF D,L-endopeptidases and to investigate the relationships between their LysM domains, by identifying orthologs of these enzymes in closely related bacterial species and comparing their LysM domains through cluster analysis.

Orthologs of CwlS, LytE and LytF were identified in 19 *Bacillus* species by the levels of amino acid identity shown by their catalytic domains (the complete list of proteins is shown in Table S1 of Supplementary Material). Results showed that all strains possessed an ortholog to LytE and LytF. However, only 13 of the 19 Bacillus species encoded a CwlS ortholog. Interestingly, the number of LysM motifs within these orthologs ranged from two to five. By these characteristics the investigated strains could be divided into groups that agreed with their respective phylogenetic lineages.

Group (A) is comprised mainly of *Bacillus licheniformis* and *Bacillus paralicheniformis* strains. Members do not possess a CwlS ortholog, but encode a LytE and a LytF orthologs with 2 and 4 LysM domains respectively. Group (B) is comprised mainly of *Bacillus amyloliquefaciens* strains, each having a CwlS ortholog with 4 LysM domains, and LytE and LytF orthologs with 2 and 3 LysM domains respectively. Group C is completely comprised of *Bacillus subtilis* strains each encoding a CwlS (4 LysM domains), LytE (3 LysM domains) and LytF orthologs (5 LysM domains). Interestingly, *Bacillus subtilis* strain XF-1 is unusual, in that it has a CwlS orthologs with 3 instead of 4 LysM domains (group X). The relationships between the 185 LysM domains in this cohort of LytE, LytF and CwlS enzymes was established based on an alignment of their amino acid sequence, followed by estimation of a phylogenetic tree with Maximum Likelihood method. The results (Figure 1), supported by high bootstrap values show that (i) individual LysM domains cluster according to their position in the protein relative to the catalytic domain; (ii) the N-terminal LysM domain is significantly diverged from those at the other positions; (iii) LysM domains group according to the enzyme type of their respective catalytic domains (i.e. LytE, LytF, CwlS) and (iv) the LysM domains of LytF and CwlS are more similar to each other than to those of LytE. The three LysM domains of the CwlS orthologue of *Bacillus subtilis* XF-1 (M4KSM2) are in that each clusters with the LysM domains at positions 2, 3, and 4 respectively, rather than with those at positions 1, 2, and 3. Since M4KSM2 is the only CwlS ortholog identified with three instead of four LysM domains, it is likely that this enzyme has undergone a deletion, losing the N-terminal LysM domain.

**Figure 1.**
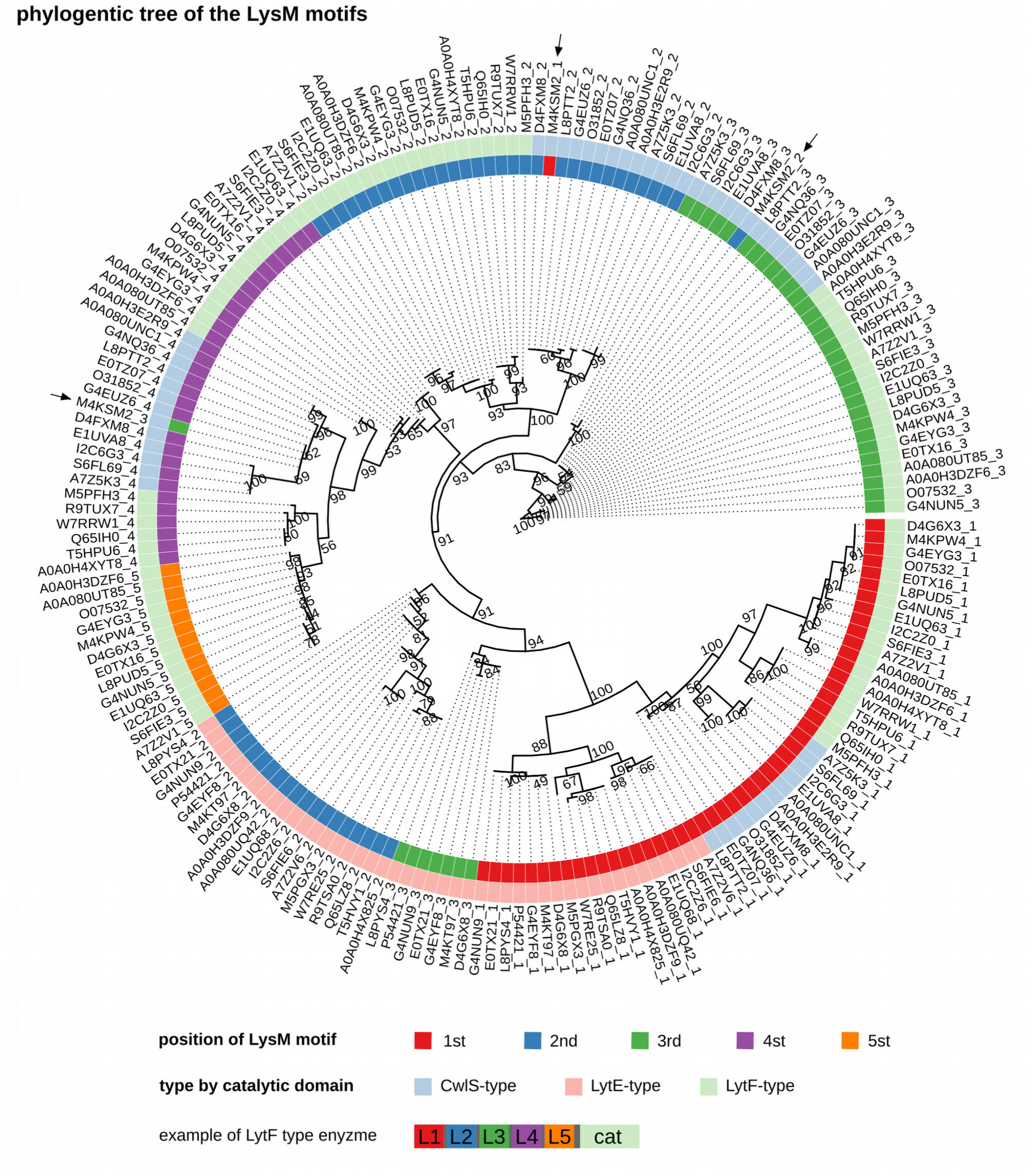
<u>Circular phylogenetic tree of the LysM motifs of the identified CwlS, LytE and LytF orthologs.</u> The amino acid sequence of the LysM motifs of the CwlS, LytE and LytF orthologs were aligned and a phylogenetic tree estimated with Maximum Likelihood method. Bootstrap values are shown (100 replicates). Inner colours correspond to the position of the individual LysM motif within the respective protein starting at the N-terminus (red= first position, blue = second position, green = third position, purple = fourth position, orange = fifth position). Outer colours correspond to the type of the respective catalytic domain (blue=CwlS, pink=LytE, green=LytF). The three LysM motifs of the CwlS-type protein M4KSM2 are highlighted by arrows. The simplified structure of a LytF-type enzyme is shown.

To further investigate LysM domain duplication, the Uniprot database was interrogated to identify the occurrence of identical LysM domains. Of ∼39,000 LysM domains identified, 46 occurred more than once, involving a total of 107 individual domains. However the duplicated domains were located in the same protein in all instances, and were adjacent to each other in 97 of the 107 instances discovered. This suggests that duplication of LysM domains occur within open reading frames with no evidence of horizontal domain transfer between proteins. These results indicate that LysM domain expansion and reduction contributes to gene divergence and that LysM domains are transferred through stable vertical inheritance.

### LytE can fulfil the D,L-endopeptidase requirement for cell viability when expressed from the *cwlO* promoter

As discovered by Bisicchia et al. (2007), LytE and CwlO are synthetically lethal, i.e. the cell requires one or other of these enzymes for viability. It is important to note that although LytE and CwlO possess similar C-terminal domains, they have different N-terminal domains and are expressed under different circumstances. The regulation of CwlO expression and activity is well described by Noone et al. (2014). In short, the TCS WalRK is the sole activator for CwlO expression, which is therefore only expressed during exponential growth, when the WalRK system is active (Salzberg *et al*., 2013). LytE expression is also activated by the WalRK system, but LytE expression can also be triggered by active SigI under stress conditions (Tseng and Shaw, 2008; Tseng *et al*., 2011; Hashimoto *et al*., 2012).

We sought to investigate whether LytE when expressed from the *cwlO* promoter can complement expression of CwlO. Since it was shown that the *cwlO* leader sequence destabilizes the *cwlO* transcript (Noone *et al*., 2014) we were also interested in seeing if the same effect applied to the *lytE* transcript.

We decided to express LytE under the control of the *cwlO* promoter in a background with a disrupted endogenous *lytE* gene [Figure 2, a].

**Figure 2.**
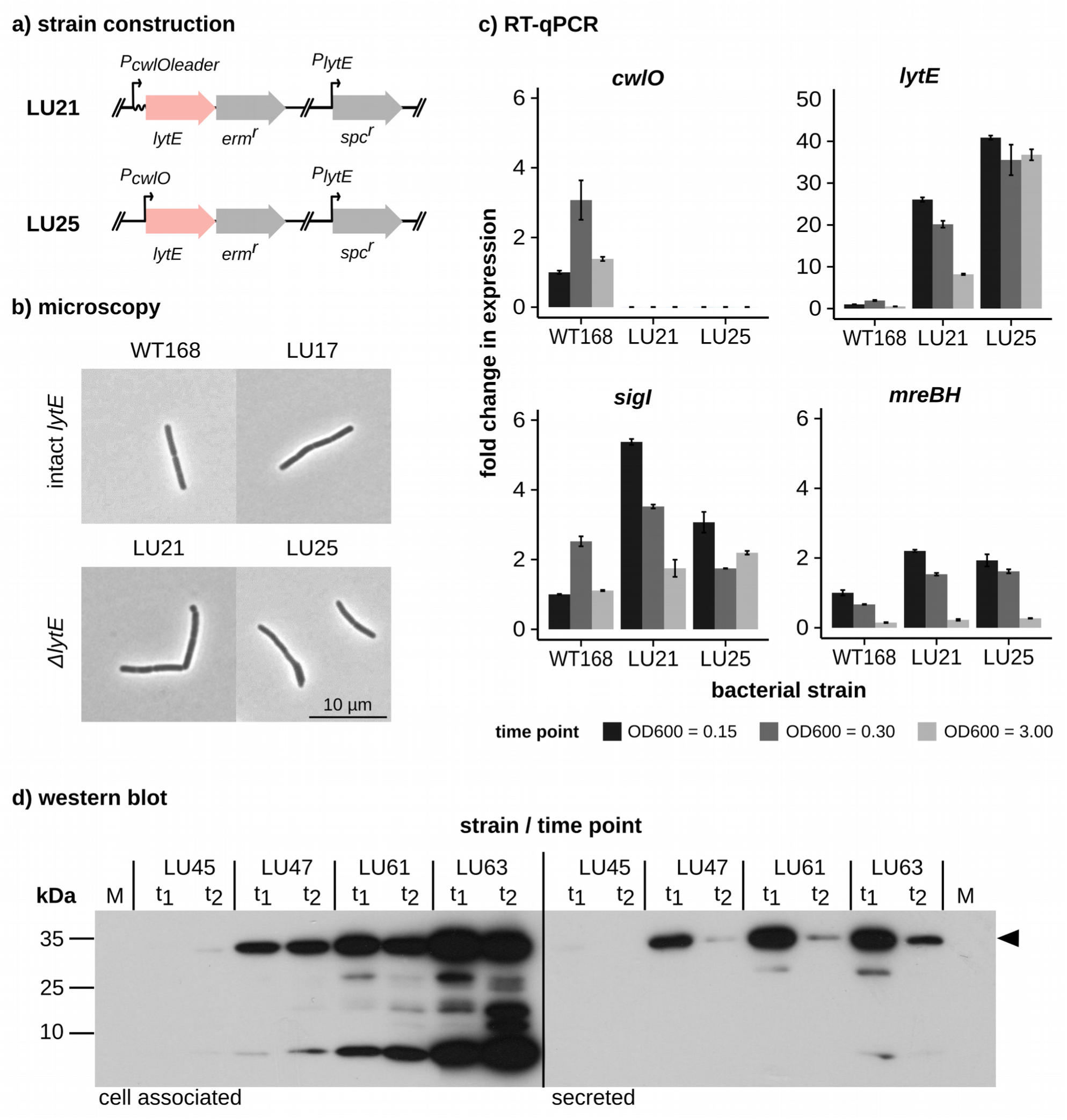
<u>2a: Schematic illustration of strains that express LytE from the *cwlO* promoter.</u> In both strains, the *cwlO* gene is replaced by the *lytE* open reading frame and the endogenous *lytE* gene is disrupted by a spectinomycin resistance cassette. In LU21 (*P_cwlOleader_lytE* Δ*cwlO* Δ*lytE*) the LytE D,L-endopeptidase is expressed from the *cwlO* promoter with *cwlO* leader sequence. In LU25 (*P_cwlO_lytE* Δ*cwlO* Δ*lytE*) the LytE enzyme is expressed from the *cwlO* promoter without *cwlO* leader sequence. Open reading frames for *lytE* and *cwlO* are not contiguous. <u>2b: Typical cell morphology of strains in which LytE is expressed from the *cwlO* promoter</u> Comparison of cell morphology of strains in which *lytE* is expressed from the *cwlO* promoter either with (LU17, LU21) or without the *cwlO* leader sequence (LU25) to the wild type (WT168). The presence or absence of the endogenous *lytE* gene is indicated. Cells were grown in LB to an OD600 of 0.1, heat-fixed and examined with an Olympus BX61 microscope. Magnification is 1000x. A scale bar is shown. <u>2c: Transcript levels of *cwlO, lytE*, *sigI* and *mreBH* in strains that express LytE from the *cwlO* promoter.</u> Transcript levels were determined by RT-qPCR for the following strains: WT168, LU21 (*P_cwlOleader_lytE* Δ*cwlO* Δ*lytE*) and LU25 (*P_cwlO_lytE* Δ*cwlO* Δ*lytE*). Cells were grown in HPDM and samples harvested at OD600=0.15, OD600=0.3 and OD600=3.00 respectively. RNA was prepared from the samples and transcribed into cDNA, which was analysed by RT-qPCR. Transcript levels were normalised relative to the level of transcript for the WT168 sample, which was assigned a value of 1. Error bars are shown. <u>2d: Cell-associated and secreted LytE protein levels in strains in which LytE is expressed from the *cwlO* promoter.</u> The level of LytE protein associated with the cell (left) and secreted in the medium (right) was established by western blot analysis in the following strains: LU45 (*lytE*-HA), LU47 (*lytE*-HA Δ*cwlO*), LU61 (*P_cwlOleader_lytE*-HA Δ*cwlO* Δ*lytE*) and LU63 (*P_cwlO_lytE*-HA Δ*cwlO* Δ*lytE*). Cells were grown in HPDM and harvested at OD600=0.2 (t1) and OD600=1.0 (t2) respectively. Total cell protein was prepared from the samples and the HA-tagged LytE protein was detected by western analysis. Twenty µg of total cell protein extract was loaded onto each lane. The amount of protein loaded for medium samples was equivalent to 1 ml medium at an OD600 = 0.3. Molecular weight markers are shown (M, kDa) and the film was exposed for 1 minute. The arrowhead indicates the position of the full-length LytE protein (LytE, MW = 37 kDa).

In comparison to the wild type, *lytE* transcript levels in such a strain were 20 to 30-fold elevated in exponentially growing cells as determined by RT-qPCR [Figure 2, c]. Furthermore a slight increase (∼ 2 to 5-fold) in *sigI* and *mreBH* transcript levels was observed. The removal of the *cwlO* leader sequence caused a further increase in *lytE* transcript and a decrease in *sigI* transcript level (∼ 2-fold). We confirmed that this can be attributed to the increased longevity of the *lytE* transcript through a mRNA half-life experiment [data not shown]. Elevated *lytE* transcript levels matched increased LytE protein levels [Figure 2, d]. High levels of LytE protein were found both cell-associated and secreted into the medium. However, the LytE protein appeared to be degraded in the medium as cultures approached stationary phase.

Although removal of the *cwlO* leader sequence led to a drop in SigI levels, we observed that overexpressing LytE at this high level affects cell morphology with a fraction of cells showing bulges and thickening [Figure 2, b]. In short, both *lytE* null mutants as well as strains overexpressing LytE, exhibit difficulties to maintain cell shape and longitudinal growth.

In summary, this results confirm that the LytE D,L-endopeptidase can complement CwlO when expressed using *cwlO* expression signals - both transcriptional (i.e. promoter) and post-transcriptional (i.e. *cwlO* leader sequence). It is clear however that the two enzymes are not equivalent since the SigI regulon (signalling cell wall stress), is induced when *lytE* is expressed using the normal *cwlO* expression signals (i.e. promoter and leader sequence), but to a lesser extent when the *cwlO* leader sequence is removed.

### LytF can also fulfil the D,L-endopeptidase requirement

The synthetic lethality of the CwlO / LytE autolysins allows further investigation of whether any other D,L-endopeptidase encoded in the genome can complement their activity. Thus, it can be investigated which D,L-endopeptidase can support growth and viability when expressed under the control of the *cwlO* promoter.

In view of these findings, we sought to establish if the D,L-endopeptidases LytF and CwlS, - which we previously showed to be closely related to LytE (LysM domains), as well as another autolytic enzyme, - YqgT (no LysM domain), can fulfil the D,L- endopeptidase requirement when expressed using the *cwlO* promoter.

To establish whether the CwlS, LytF or YqgT D,L-endopeptidases when expressed from the *cwlO* promoter can fulfill the D,L-endopeptidase requirement, we constructed strains in which the *cwlS, lytF* and *yqgT* genes were individually expressed at the *cwlO* locus.

The level of transcript of each gene was varied by expressing each gene both with the *cwlO* leader sequence (giving an unstable transcript) and without the leader sequence (giving a stable transcript). The ability of each enzyme to fulfil the D,L- endopeptidase requirement for viability was then established by attempting to disrupt the *lytE* gene in each of the strains, which was successful for *lytF* but not for *cwlS* or *yqgT*. LytF levels in the strains expressing LytF from the *cwlO* promoter were highly elevated when compared to that observed in strains, in which the enzyme was expressed from its natural promoter [Figure 3, b]. Despite the high level of cellular *lytF* mRNA transcript, only a very low level of full-length LytF protein was cell associated, with a high fraction found to be processed [Figure 3, c]. HA-tagged LytF protein was predominantly localized at the cell septa and cell poles, but was also present along the lateral cell cylinder [Figure 3, d]. Based on the *sigI* transcript levels, it seems that significant cell wall stress is not generated in these strains [data not shown]. However, strains overexpressing LytF from the *cwlO* promoter displayed major growth defects (delayed lag phase) and cell morphology defects [Figure 3, a].

**Figure 3.**
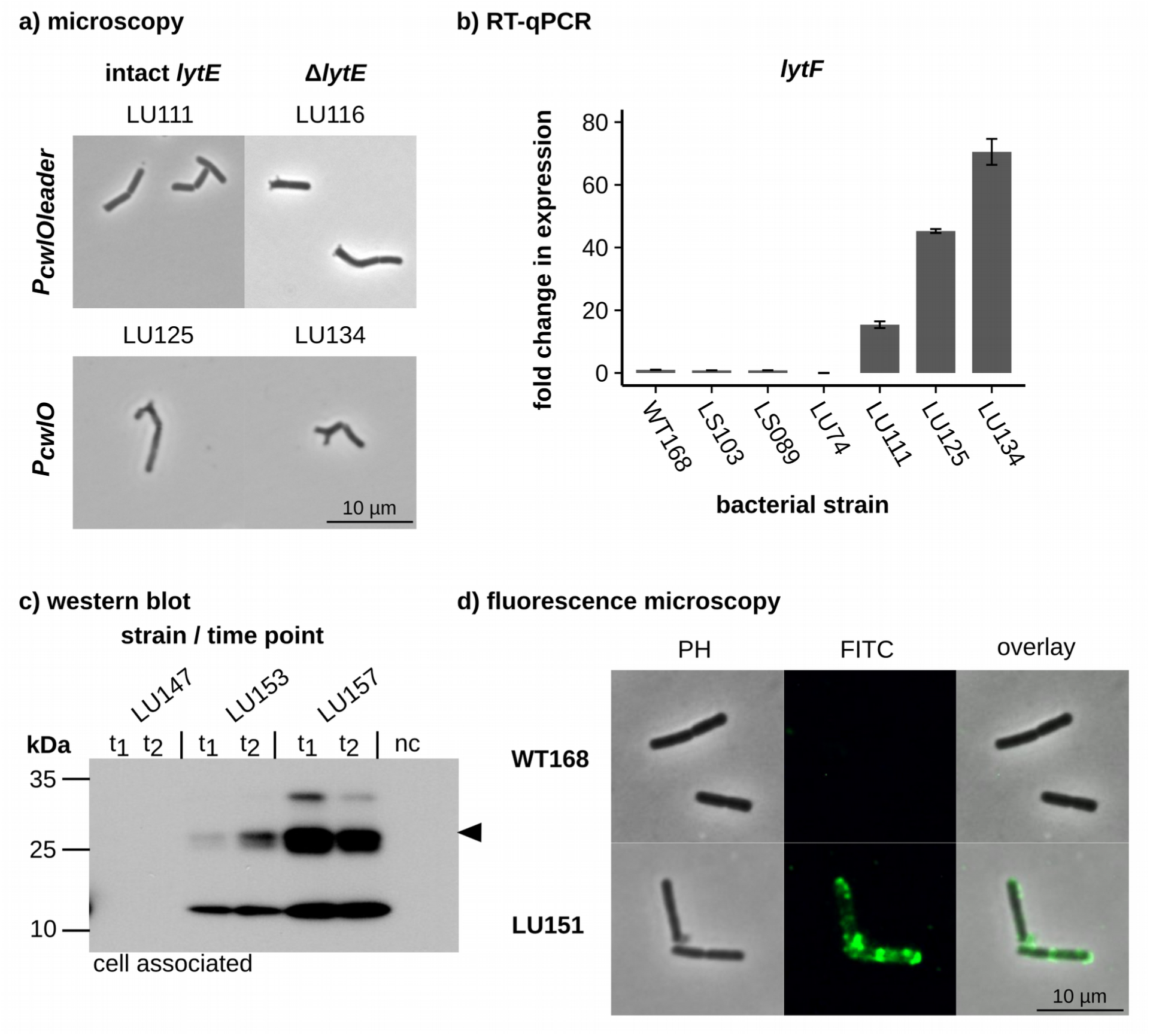
<u>3a: Typical cell morphology of strains that express LytF from the *cwlO* promoter.</u> Comparison of cell morphology of strains in which *lytF* is expressed from the *cwlO* promoter either with (LU111, LU116) or without the *cwlO* leader sequence (LU125, LU134). The presence or absence of the endogenous *lytE* gene is indicated. The endogenous *cwlO* and *lytF* genes are disrupted in all strains. Cells were grown in LB to an OD600 of 0.1, heat-fixed and examined with an Olympus BX61 microscope. Magnification is 1000x. A scale bar is shown. <u>3b: Transcript levels of *lytF*.</u> LytF transcript levels were determined by RT-qPCR, for the following strains: WT168, LS103 (Δ*cwlO*), LS089 (Δ*lytE*), LU74 (Δ*lytF*), LU111 (*P_cwlOleader_lytF* Δ*lytF* Δ*cwlO*), LU125 (*P_cwlO_lytF* Δ*lytF* Δ*cwlO*) and LU134 (*P_cwlO_lytF* Δ*lytF* Δ*cwlO* Δ*lytE*). Cells were grown in HPDM and samples harvested at OD600=0.15. RNA was prepared from the samples and transcribed into cDNA, which was analysed by RT-qPCR. Transcript levels were normalised relative to the level of transcript for the WT168 sample, which was assigned a value of 1. Error bars are shown. <u>3c: Cell-associated protein levels of LytF in WT168 and when expressed from the *cwlO* promoter with and without *cwlO* leader sequence.</u> Cell-associated protein samples of the following strains are shown: LU147 (*lytF*-HA), LU153 (Δ*lytF P_cwlOleader_lytF*-HA Δ*cwlO* Δ*lytE*) and LU157 (Δ*lytF P_cwlO_lytF*-HA Δ*cwlO* Δ*lytE*). A negative control (WT168 without HA-tag) is also displayed. Cells were grown in HPDM and harvested at OD600=0.2 (t1) and OD600=1.0 (t2). Total cell protein was prepared from the samples and the HA-tagged proteins were detected by western analysis. Ten µg total cell protein extract was deployed per lane. Molecular weight markers are shown (M, kDa). The film was exposed for one minute. The arrowhead indicates the degraded LytF protein (full length LytF, MW = 51 kDa). <u>3d: Localization of the LytF protein in a strain in which the protein is expressed from the *cwlO* promoter.</u> Strain LU151 (Δ*lytF P_cwlOleader_lytF*-HA Δ*cwlO*) was grown in buffered LB to an OD600 of 0.2 at which point cells were harvested, fixed in 2.6 % paraformaldehyde, immunostained and examined with an Olympus BX61 microscope. Immunostaining was achieved by endowing the LytF protein with a HA-tag and treating cells first with a primary anti-HA antibody followed by binding of the primary antibody with a fluorescein isothiocyanate (FITC)-labelled secondary antibody. Proteins were visualized by phase contrast microscopy (PH) and by immunofluorescence microscopy (FITC). Magnification is 1000x. A negative control (nc) is displayed (WT168, no HA-tag). A scale bar is shown.

Thus, while cell viability can be achieved by replacing the CwlO and LytE D,L- endopeptidases with expression of LytF, cells are not robust.

### The individual LysM domains of LytE are not functionally equivalent

We could show that LytE (3 LysM domains) and LytF (5 LysM domains), but not CwlS (4 LysM domains) can support growth of *B. subtilis*, when expressed from the *cwlO* promoter. Interestingly although the majority of LysM domain containing proteins in the dataset constructed in section one possess only a single LysM domains (∼ 67 % of 18,246 proteins), all identified orthologs to LytE, CwlS and LytF feature at least 2 LysM domains.

There is little known about how individual LysM domains bind to peptidoglycan and the significance of the fact that the identified D,L-endopeptidase orthologs possess a minimum of 2 LysM domains.

Therefore we sought to investigate the properties of the individual LysM domains of LytE by designing LytE enzymes with a single LysM domain. We then tested the characteristics of these LytE variants proteins in strains which viability was depended on the permutated autolysin. This was achieved by expressing the variant LytE containing either L1, L2 or L3 in single copy at the *cwlO* or the *lytE* locus and attempting to disrupt either the endogenous *lytE* or the *cwlO* gene [Supplementary Figure 2]. For control purposes, we also included a variant LytE without any LysM domain as well as a reconstituted wild type LytE with the three LysM domains in the wild type combination (L1-L2-L3) in the further analysis.

We found that all constructed strains were viable, demonstrating that any single LysM domain of LytE is capable of inferring functionality to the LytE enzyme. A control strain with a LytE protein without a LysM domain could not be constructed, demonstrating that the LytE enzyme requires at least one LysM domain for functionality.

Despite the fact that any individual LysM domain enabled a functional LytE enzyme, we detected a significant change in cell morphology with shorter and thicker cells for the variant strains. Interestingly this effect was greater for strains containing a LytE enzyme with L2 or L3 than L1 [Figure 4, a].

**Figure 4.**
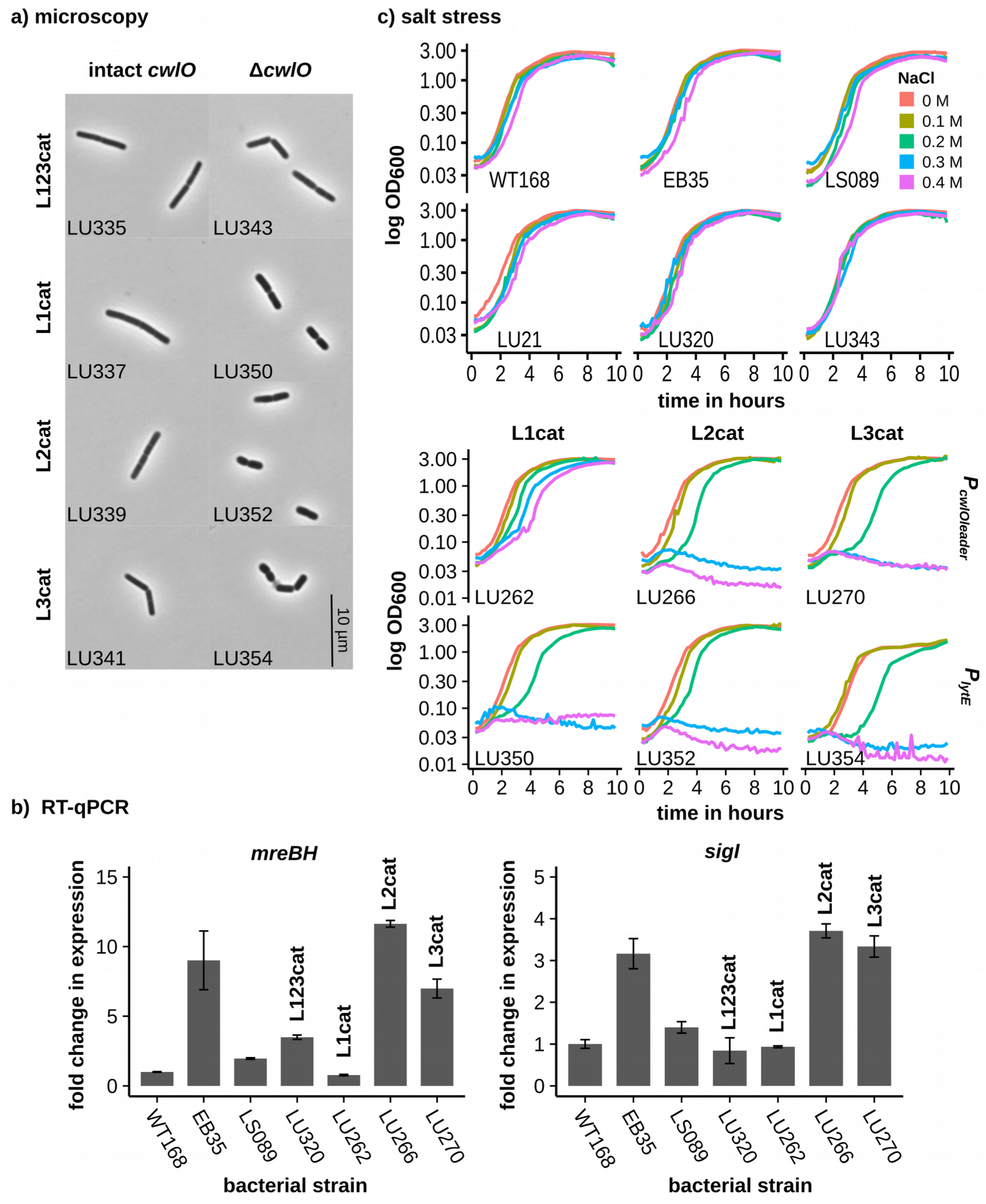
<u>4a: Typical cell morphology of strains that express a variant LytE protein with one LysM motif from the *lytE* promoter either with or without the endogenous *cwlO* gene.</u> Comparison of cell morphology of strains in which a variant LytE with one LysM motifs is expressed from the *lytE* promoter in combination with an intact *cwlO* gene (LU337 (*P_lytE_lytE[L1cat] ΔlytE*), LU339 (*P_lytE_lytE[L2cat] ΔlytE*), LU341 (*P_lytE_lytE[L3cat] ΔlytE*)), or in combination with a *cwlO* null mutant (LU350 (*P_lytE_lytE[L1cat] ΔlytE ΔcwlO*), LU352 (*P_lytE_lytE[L2cat] ΔlytE ΔcwlO*), LU354 (*P_lytE_lytE[L3cat] ΔlytE ΔcwlO*)). A control with a reconstituted wild type *lytE* is shown (LU335 (*P_lytE_lytE[L123cat]R ΔlytE*), LU343 (*P_lytE_lytE[L123cat]R ΔlytE ΔcwlO*)). Cells were grown in LB to an OD600 of 0.1, heat-fixed and examined with an Olympus BX61 microscope. Magnification is 1000x. A scale bar is shown. <u>4b: *sigI* and *mreBH* transcript levels for strains that express a variant LytE protein from the *cwlO* promoter.</u> Transcript levels were established by RT-qPCR for the following strains: WT168, EB35 (Δ*cwlO*), LS089 (Δ*lytE*), LU320 (*P_cwlOleader_lytE[L123cat]R* Δ*cwlO* Δ*lytE*), LU262 (*P_cwlOleader_lytE[L1cat]* Δ*cwlO* Δ*lytE*), LU266 (*P_cwlOleader_lytE[L2cat]* Δ*cwlO* Δ*lytE*) and LU270 (*P_cwlOleader_lytE[L3cat]* Δ*cwlO* Δ*lytE*). Cells were grown in HPDM and samples harvested at OD600=0.15. RNA was prepared from the samples and transcribed into cDNA, which was analysed by RT-qPCR with specific primer sets. Transcript levels were normalised relative to the level of transcript for the WT168 sample, which was assigned a value of 1. Error bars are shown. <u>4c: Growth curves of strains that express a variant LytE protein with one LysM motif from the *cwlO* and the *lytE* promoter at different salt concentrations.</u> Growth curves were obtained for the following strains: WT168, EB35 (Δ*cwlO*), LS089 (Δ*lytE*), LU21 (*P_cwlOleader_lytE* Δ*cwlO* Δ*lytE*), LU320 (*P_cwlOleader_lytE[L123cat]R* Δ*cwlO* Δ*lytE*), LU343 (*P_lytE_lytE[L123cat]R* Δ*cwlO* Δ*lytE*), LU262 (*P_cwlOleader_lytE[L1cat]* Δ*cwlO* Δ*lytE*), LU266 (*P_cwlOleader_lytE[L2cat]* Δ*cwlO* Δ*lytE*), LU270 (*P_cwlOleader_lytE[L3cat]* Δ*cwlO* Δ*lytE*), LU350 (*P_lytE_lytE[L1cat]* Δ*cwlO* Δ*lytE*), LU352 (*P_lytE_lytE[L2cat]* Δ*cwlO* Δ*lytE*), LU354 (*P_lytE_lytE[L3cat]* Δ*cwlO* Δ*lytE*). Growth curves were obtained in the following way: Overnight cultures of the strains were grown in salt-free buffered LB and used to inoculate 100 µl media in a CELLSTAR® 96 well plate to an OD600 of 0.03. NaCl was added to the final concentrations of 0.1 M, 0.2 M, 0.3 M and 0.4 M. The 96 well plate was incubated at 37°C with slow shaking and the OD600 was measured every 10 minutes over a time period of 10 hours. Three technical repeats for each strain were conducted. One representative example is shown.

We suggest that the extent to which a variant LytE protein fulfils the D,L-endopeptidase requirement can be assessed by determining the extent to which the SigI regulon is induced. This was determined by establishing the mRNA transcript levels of the SigI regulon genes *sigI* and *mreBH* by RT-qPCR. We observed that transcript levels were elevated if a strain contained a variant LytE with a single LysM domain with the combination L2, L3, however no elevation occurred with L1 or with the reconstituted wild type protein [Figure 4, b].

We further decided to test the robustness of these strains by determining their sensitivity to salt stress. As expected, all controls grew normally at NaCl concentrations up to 0.4 M. In contrast, although all variant LytE strains grew comparable to wild type under the absence of salt or at a salt concentration of 0.1 M, growth inhibitions were observed at a salt concentration of 0.2 M and greater [Figure 4, c]. The extent, to which salt stress caused a change in growth behaviour, correlated with the type of LysM domains as well as the expression level (*cwlO* vs *lytE* promoter). Higher expression levels (*cwlO* promoter) improved the resistance of strains to salt stress. Salt tolerance was also increased when the LytE enzyme contained the first LysM domain (L1) as compared to L2 and L3. Since sensitivity to salt stress was not altered in the control strains which were in possession of an intact copy of the endogenous *lytE* or *cwlO* gene, we conclude that it is not the presence of the variant LytE that causes the increased sensitivity to salt stress, but rather a lack of D,L-endopeptidase enzymatic activity.

Our results indicate that although a variant LytE protein with any individual LysM domain is functional, it does not fulfil the D,L-endopeptidase requirement to the same extent as the wild type LytE protein. Cells that express a variant LytE with a single LysM domain, under the absence of the endogenous *cwlO* and *lytE* gene display growth and morphology defects. Furthermore our experiments show that the three LysM domains of LytE are not functionally equivalent and that L1 is capable of conferring on the catalytic domains the ability to fulfil the D,L-endopeptidase requirement to a greater extent than L2 and L3 as can be seen from the induction of the *sigI* regulon.

### LysM domain requirement of LytE

We observed that the three LysM domains are not equal in their ability to confer functionality to the LytE enzyme with the N-terminally located LysM domain L1 being superior to L2 and L3. It is not clear what properties distinguish the first LysM domain from the subsequent LysM domains, other than its location adjacent to the signal sequence domain.

We therefore sought to investigate the effect of the order as well as the number of LysM domains on the functionality of the LytE enzyme.

This was achieved by expressing variant LytE enzymes with two or three LysM domains in a permutated order from either the *cwlO* or the *lytE* promoter [Supplementary Figure 2]. Functionality of these enzymes was then tested by disrupting the endogenous *lytE* or *cwlO* gene. We could show that any LytE variant with two LysM domains in any of the nine possible combinations resulted in a functional enzyme. The same was true for two variant LytE with three LysM domains in the combinations L3-L3-L1 and L2-L1-L3.

Although all strains were viable, we observed that strains expressing variant LytE proteins with three LysM domains produced bigger colonies than those that expressed variant LytE proteins with two LysM domains [Figure 5, a]. Strains containing a variant

**Figure 5.**
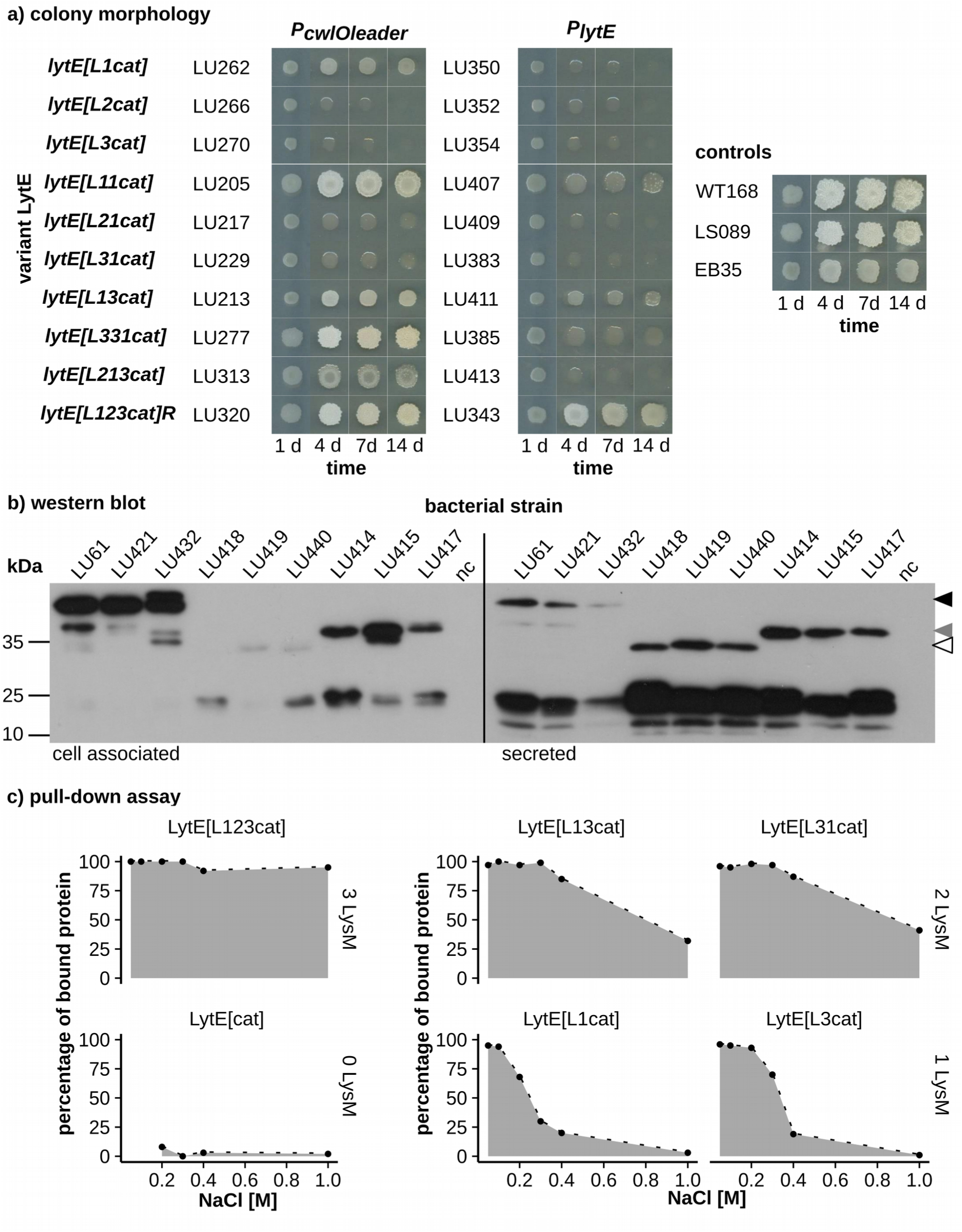
<u>5a: Colony morphology of strains that express a variant LytE protein from either the *cwlO* or the *lytE* promoter.</u> LB agar plates (no antibiotic) were inoculated with a toothpick from an overnight LB culture and incubated overnight at 37°C followed by incubation for 14 days at room temperature. Growth of each colony was documented after 1 day, 4 days, 7 days and 14 days. The following strains were characterized: LU262 (*P_cwlOleader_lytE[L1cat]* Δ*cwlO* Δ*lytE*), LU266 (*P_cwlOleader_lytE[L2cat]* Δ*cwlO* Δ*lytE*), LU270 (*P_cwlOleader_lytE[L3cat]* Δ*cwlO* Δ*lytE*), LU205 (*P_cwlOleader_lytE[L11cat]* Δ*cwlO* Δ*lytE*), LU217 (*P_cwlOleader_lytE[L21cat]* Δ*cwlO* Δ*lytE*), LU229 (*P_cwlOleader_lytE[L31cat]* Δ*cwlO* Δ*lytE*), LU213 (*P_cwlOleader_lytE[L13cat]* Δ*cwlO* Δ*lytE*), LU277 (*P_cwlOleader_lytE[L331cat]* Δ*cwlO* Δ*lytE*), LU313 (*P_cwlOleader_lytE[L213cat]* Δ*cwlO* Δ*lytE*), LU320 (*P_cwlOleader_lytE[L123cat]R* Δ*cwlO* Δ*lytE*), LU350 (*P_lytE_lytE[L1cat]* Δ*lytE* Δ*cwlO*), LU352 (*P_lytE_lytE[L2cat]* Δ*lytE* Δ*cwlO*), LU354 (*P_lytE_lytE[L3cat]* Δ*lytE* Δ*cwlO*), LU407 (*P_lytE_lytE[L11cat]* Δ*lytE* Δ*cwlO*), LU409 (*P_lytE_lytE[L21cat]* Δ*lytE* Δ*cwlO*), LU383 (*P_lytE_lytE[L31cat]* Δ*lytE* Δ*cwlO*), LU411 (*P_lytE_lytE[L13cat]* Δ*lytE* Δ*cwlO*), LU385 (*P_lytE_lytE[L331cat]* Δ*lytE* Δ*cwlO*), LU413 (*P_lytE_lytE[L213cat]* Δ*lytE* Δ*cwlO*), LU343 (*P_lytE_lytE[L123cat]R* Δ*lytE* Δ*cwlO*), WT168, LS089 (Δ*lytE*), EB35 (Δ*cwlO*). <u>5b: The levels of LytE variant proteins expressed from the *cwlO* promoter, for cell-associated proteins and secreted proteins.</u> Protein samples of the following strains are shown: LU61 (Δ*cwlO P_cwlOleader_lytE*-HA Δ*lytE*), LU421 (*P_cwlOleader_lytE*-HA *[L123cat]R* Δ*cwlO* Δ*lytE*), LU432 (*P_cwlOleader_lytE*-HA *[L213cat]* Δ*cwlO* Δ*lytE*), LU418 (*P_cwlOleader_lytE*-HA *[L1cat]* Δ*cwlO* Δ*lytE*), LU419 (*P_cwlOleader_lytE*-HA *[L2cat]* Δ*cwlO* Δ*lytE*), LU440 (*P_cwlOleader_lytE*-HA *[L3cat]* Δ*cwlO* Δ*lytE*), LU414 (*P_cwlOleader_lytE*-HA *[L11cat]* Δ*cwlO* Δ*lytE*), LU415 (*P_cwlOleader_lytE*-HA *[L13cat]* Δ*cwlO* Δ*lytE*) and LU417 (*P_cwlOleader_lytE*-HA *[L31cat]* Δ*cwlO* Δ*lytE*). A negative control nc (WT168 without HA-tag) is also displayed. Cells were grown in salt-free buffered LB and harvested at an OD600 of 0.25. Total cell protein was prepared from the samples and the HA-tagged proteins were detected by western analysis. Five µg total cell protein extract was deployed per lane (left) and 0.15 OD equivalent media proteins were deployed per lane (right). Molecular weight markers are shown (M, kDa). Both films were exposed for seven minutes. The black arrowhead indicates the full length LytE protein (LytE, MW = 37 kDa), the grey arrowhead indicates a full length dimer protein (variant LytE with two LysM motifs, MW = 30 kDa) and the open arrowhead indicates a full length monomer protein (variant LytE with one LysM motif, MW = 23 kDa). <u>5c: *In vitro* binding affinity of purified variant LytE proteins to purified *B. subtilis* cell walls at different salt concentrations.</u> The diagram shows the results of peptidoglycan pull down assays, conducted for purified variant LytE proteins by proportion of protein found in the pellet. The following proteins were analysed: LytE[L123cat], LytE[cat], LytE[L13cat], LytE[L31cat], LytE[L1cat], LytE[L3cat]. Forty-four pmole of variant LytE protein were mixed with purified cell wall of *B. subtilis* to a final absorbance of OD600 = 10, in a 25 µl reaction volume. A negative control (nc) with without cell wall was performed to ensure that proteins do not precipitate on their own. Pull-down assays were conducted at five different NaCl concentrations: 0.05 M, 0.1 M, 0.2 M, 0.3 M, 0.4 M and 1 M.

LytE protein with a single LysM domain displayed the smallest colonies, which also exhibited strong lysis over the course of the experiment. In general, colonies were bigger and experienced less lysis when they were expressed using the stronger *cwlO* promoter rather than the weaker *lytE* promoter. Interestingly the order of LysM domains also mattered. Those strains that expressed a variant LytE with L1 at the first (N-terminal) position displayed larger colonies, than those in which either L2 or L3 were located adjacent to the signal sequence domain. This correlation was especially strong for LytE variants with a set of 2 LysM domains, as can be seen when comparing L1-L3 to L3-L1.

To address whether the type of variant LytE protein has an effect on the cellular protein level, the level of HA-tagged variant LytE protein was determined by western blot analysis. All variant LytE-HA proteins could be detected and bands migrated according to the predicted MW [Figure 5, b]. The level of variant LytE protein varied significantly in the strains tested. Since the number of LysM domains is proposed to correlate with binding affinity, we expected to find more cell-associated protein in variant proteins with a greater number of LysM domains. Indeed, the level of cell-associated variant LytE protein was higher for proteins with two or more LysM domains, than for proteins with a single LysM domain. If it is assumed that the lower level of cell-associated LytE is caused by lowered binding to the cell wall, and not lowered expression, then we predict the level of variant LytE protein found in the medium to have the reverse profile of that for cell-associated protein. To verify this, we compared the levels of cell-associated variant LytE proteins with those in the culture supernatant. A strong inverse correlation in the level of variant LytE was observed. Variant LytE proteins with one LysM domain were predominantly found in the culture supernatant, while variant LytE proteins with three LysM domains were predominantly found in association with the cell wall. Moreover the distribution of variant LytE proteins with two LysM domains takes an intermediate between that found for those with three and one LysM domains. We confirmed these results by immunofluorescence microcopy, which demonstrated that HA tagged variant LytE with more LysM domains gave a stronger fluorescence signal than those with fewer LysM domains [data not shown]. Interestingly, the pattern of localization along the cell cylinder was not affected.

We further tested binding affinity of variant LytE protein in vitro in an insoluble peptidoglycan pull down assay. To investigate the effect of salt on the peptidoglycan binding affinity we added increasing amounts of NaCl to the assay buffer (0.05 M - 1 M NaCl) [Figure 5, c]. At a low salt concentration of 0.05 M NaCl, all variant LytE proteins showed an equally strong binding capacity of approximately 100 % (proportion of protein found in the pellet). However, no binding could be detected for the variant LytE protein without LysM domain. Furthermore our results clearly show that the peptidoglycan-binding capacity is negatively correlated to the salt concentration. With an increase in salt concentration (≥ 0.1 M NaCl) binding capacity for the variant LytE proteins is decreased. The rate at which binding capacity is declined varies between proteins and is in negative correlation to the number of LysM domains present. Proteins with more LysM domains exhibited a greater robustness of binding affinity than proteins with a single LysM domain.

In summary, this study demonstrates that the number and order of LysM domains are of great relevance for the functionality of the LytE enzyme. Although a LytE enzyme with any one of its three LysM domains is functional and can fulfil the D,L-endopeptidase requirement, increasing the number of LysM domains to two or three has a positive effect on normal colony morphology. The order in which the domains are arranged is also of importance in that variant LytE proteins in which the first LysM domain (L1) is localized in its natural position adjacent to the signal peptide fulfil the D,L-endopeptidase requirement to a greater extent than L2 and L3.

Furthermore, the number of LysM domains of a LytE variant protein is positively correlated with peptidoglycan binding affinity and a good determinant of the extent to which the protein is cell-associated or shed into the culture media. We could show that an increase of the number of LysM domains leads to increased binding affinity both *in vivo* and *in vitro*. We further demonstrated that binding affinity is negatively affected by salt *in vitro*.

## DISCUSSION

### The WalRK and SigIRsgI systems work in tandem to ensure adequate expression of D,L-endopeptidase activity

Expression of D,L-endopeptidase activity is required for normal growth and viability of *B. subtilis* as reported by Bisicchia et al. (2007). Under normal growth condition this activity is provided by CwlO and under stress conditions by the LytE enzyme. We demonstrated in this study that the level of *lytE* transcript and LytE protein is significantly increased when the *cwlO* gene is deleted, probably to compensate for the loss in D,L- endopeptidase activity. Since *lytE* is part of the SigI regulon (Tseng *et al*., 2011), we suggest that this is achieved through activation of the alternative SigI factor as proposed by Salzberg and colleagues (2013). A similar response was observed by Kasahara and colleagues (2016) who reported that *lytE* transcription was considerably increased in strains with defects in teichoic acid synthesis, notably in a *ltaS* null mutant and who suggest that upregulation is achieved in a SigI dependent manner. Thus, the WalRK and SigIRsgI regulators form a regulatory network that functions to ensure that there is an adequate level of D,L-endopeptidase enzyme present under normal growth (CwlO) and cell wall stress (LytE) conditions.

It was of interest to establish whether LytE can fulfil the D,L-endopeptidase requirement for viability when expressed from the *cwlO* promoter in the absence of the endogenous *lytE* gene (thus removing SigI coupling). It was found that such a strain is viable, signifying that changing the regulation of LytE expression from its natural promoter to the *cwlO* promoter is compatible with viability and indeed normal growth. We observed that LytE expression was higher when driven by the *cwlO* promoter as compared to the endogenous *lytE* promoter (un-induced or induced). LytE expression (stabilized transcript and increased protein level) was further increased by deletion of the *cwlO* leader sequence due to the destabilizing effect of the leader transcript (Noone *et al*., 2014). Surprisingly, we detected a slight increase in *mreBH* and *sigI* when *lytE* was expressed from the *cwlO* promoter with the *cwlO* leader sequence. Based on the premise that *sigI* is released under stress, removal of the *cwlO* leader sequence is beneficial for the cell, since it results in a decrease of *sigI* levels. However, at the same time removal of the *cwlO* leader sequence caused minor deficiencies in cell morphology, indicating that such an elevated level of autolytic activity is detrimental to the cell. The time point at which these phenotypic defects were observed, correlates with the high level of WalRK activity during these growth phases (Prunty *et al*., 2018)

In summary, these data suggest that although expression of LytE from the *cwlO* promoter can fulfil the D,L-endopeptidase requirement for cell viability, it does not complement CwlO function completely and that further elevation of LytE protein levels are required (i.e. by removal of the leader sequence) to achieve a state without a cell wall stress (i.e. no elevation of *sigI* expression).

### Expression levels and localization of D,L-endopeptidases within the cell envelope are critical

We investigated four of the seven D,L-endopeptidases encoded in the *B. subtilis* genome (LytE, LytF, CwlS, YqgT) and found that CwlS and YqgT cannot complement CwlO / LytE activity when expressed from the *cwlO* promoter. LytE naturally fulfils the D,L-endopeptidase requirement in a *cwlO* mutant, but can also do so when expressed from the *cwlO* locus. LytF cannot complement CwlO / LytE activity when expressed from its natural *lytF* promoter, but gives partial complementation when expressed from the *cwlO* promoter. The fact that LytF is unable to fulfil the D,L-endopeptidase requirement for viability when expressed from the *lytF* promoter, but can do so when expressed from the *cwlO* promoter, indicates that complementation requires either greater than normal LytF protein levels and / or LytF expression at a different time point throughout the growth cycle. However, although a viable strain could be constructed, severe growth and morphology defects were observed. Exacerbation of the aberrant phenotype in strains without *cwlO* leader sequence (causing a further increase in LytF levels) suggests that the level of LytF in this background may be too high for normal growth.

We do not know why neither *cwlS* nor *yqgT* were able to fulfil the D,L-endopeptidase requirement. However, we ruled out the possibility of a synthetic lethality between these enzymes by constructing viable double mutants.

The fact that the LytE, LytF and CwlS enzymes are highly similar in their C-terminal D,L- endopeptidase catalytic domains, as well as in their N-terminally located LysM domains poses the question as to what causes the inability of CwlS to fulfil the D,L- endopeptidase requirement. We confirmed that there is indeed a high level of CwlS expressed and associated with the cell, when expressed from the *cwlO* promoter [data not shown]. However, it may be of relevance that the CwlS protein is primarily located at the cell septa (Hashimoto *et al*., 2012), while both LytE and LytF have some presence along the cell cylinder (Hashimoto *et al*., 2012; Kiriyama *et al*., 2014; Kasahara *et al*., 2016). This is significant since it has been established that the D,L-endopeptidase requirement is associated with facilitating cell cylinder elongation (Bisicchia *et al*., 2007). Thus, there may not be a sufficient level of CwlS located along the cylinder to facilitate cell elongation. This is supported by the observations that cells expressing *cwlS* from the *cwlO* promoter appear to be shorter than wild-type cells and to have unusual structures at the poles of the cell (data not shown).

Although both LytE and LytF show some binding to the lateral cell wall, LytF is primarily bound to the cell septa and poles, which potentially explains why LytF achieves only partial complementation in comparison to the LytE enzyme. In summary, these results show that beside the level of D,L-endopeptidase expression, their proper subcellular localization are paramount for normal growth.

### The individual LysM domains of LytE are functionally and structurally distinct

We could show that strains expressing a LytE variants with any single LysM domain are viable. However, we observed cell morphology, colony morphology and growth defects in such strains. These phenotypic defects were more pronounced in LytE variants with a single copy of the second (L2) or third (L3) LysM domains as opposed to the N-terminal located L1, indicating that the individual LysM domains are not functionally equivalent. The increased effectiveness of the first LysM domain (L1) is probably due to the fact that it has evolved to be more suitable for interaction with the adjacent signal sequence domain, which perhaps causes the enzyme to be expressed or exported more efficiently.

This finding was confirmed in later experiments when we could show that in strains that express a variant LytE proteins with two or three LysM domains, the order of the domains was relevant for proper colony growth. Variant LytE proteins that contained the first LysM domain (L1) at the N-terminal position resulted in strains that displayed healthier colonies which experienced less lysis than variant LytE in which either L2 or L3 were located adjacent to the signal peptide. A possible mechanism by which a LytE protein with impaired functionality could cause colony growth defects was investigated by Mamou and colleagues (2017), who report that LytE is involved in proper colony development through influencing Y arm extension and colony thickening patterns.

The fact that the first LysM domain is structurally different from the subsequent domains was further supported through bioinformatic analysis. By comparing the amino acid sequence of the 185 LysM domains contained in the LytE, CwlS and LytF orthologs of 19 closely related *B. subtilis* species we found that the N-terminal located LysM domains were the most distinct. This may indicate that due its interaction with the signal peptide there is a greater selection pressure on the N-terminal LysM domain than on subsequent domains.

### LysM domains contribute to peptidoglycan binding in vivo and in vitro

We could show that a variant LytE without LysM domains exhibited no binding affinity to purified cell wall *in vitro* and that it could not sustain cell growth. Thus, LysM domains are essential for cell wall binding and therefore functionality of the LytE enzyme. This observation is supported by several previous studies that have shown that a total loss of LysM domains greatly decreases the peptidoglycan-binding capacity in vitro (Steen *et al*., 2005; Eckert *et al*., 2006; Wong *et al*., 2014). Increasing the number of LysM domains from one to three, led to increased binding capacity both *in vivo* and *in vitro*, suggesting that individual LysM domains contribute to binding in a accumulative manner, as was proposed previously (Mesnage *et al*., 2014; Wong *et al*., 2014).

Surprisingly, strains expressing a LytE variant with any single LysM domain are viable, suggesting that the peptidoglycan-binding affinity of the wild type LytE protein with three LysM domains is excessive. However, our experiments were performed under laboratory conditions and we observed that such strains have elevated stress sensitivity, especially to salt stress. We demonstrated *in vitro* that salt causes a decrease in binding capacity of variant LytE proteins, especially those with fewer than three LysM domains. Therefore, growth defects in the presence of salt are probably due to insufficient binding of the variant LytE D,L-endopeptidase under this circumstances. Given that *B. subtilis* is a soil-dwelling organism, its capability to withstand salt stress is essential for its survival. Therefore, cells that possess a LytE enzyme with less than three LysM domains would not be able to survive natural fluctuations in the salt concentrations in the soil.

### Expansion of the LysM domains and evolution of the CwlS, LytE and LytF orthologs

We observed that the LysM domains of the CwlS, LytE and LytF orthologs cluster primarily by position, with the N-terminally located LysM domains forming a remarkably distinct cluster. This finding is contrary to a similar study conducted for fungal proteins. In this study, Cen and colleagues (2017) report that the LysM domains of fungal effectors proteins cluster independently of their sequential position within the parental protein or protein structure family. We suggest that this difference stems from the fact that the LytE, CwlS and LytF proteins have evolved fairly recent, leaving their LysM domains little time to diverge.

We were further interested in the manner in which D,L-endopetidases acquire new LysM domains. By screening 26.078 bacterial proteins for identical LysM domain amino acid sequences we could show that LysM domains are primarily propagated through duplication events that occur within a gene. Interestingly, the majority of duplicated LysM domains were present within the phylum Firmicutes (79.4 %). This is a much higher percentage than could be expected, provided that domain expansion occurred in all phyla at the same rate (LysM domains of the Firmicutes phylum represent 32.8 % of the total dataset), suggesting that LysM domain expansion is either a more recent or more frequent event in species of the Firmicutes phylum. Interestingly, an uncharacterised protein of *Bacillus subtilis* subsp. *natto* BEST195 (D4G4L0) was found to possess 6 LysM domains, that are of identical amino acid sequence and highly similar (although not identical) on nucleotide level. This indicates that LysM domain duplication events can create several repeats of a single domain at once, leading to the rapid expansion of the domain repeats.

Based on these findings, we propose a model for the evolutionary development of the three D,L-endopeptidases and their LysM domains. Our model is based on the premise that LytE, CwlS and LytF share a common ancestor and that LysM domain repeats expand independently within genes and are passed on through stable vertical inheritance. In this model an ancestral gene with a single N-terminally located LysM domain, undergoes a combination of gene and LysM domain duplication / loss events, which lead to the development of the three orthologs and their observed LysM domain combinations [Figure 6, a]. This model can also be used to infer basic phylogeny between the investigated *Bacillus* species [Figure 6, b], which is in line with their overall phylogenetic relationships [Supplementary Figure 1].

**Figure 6.**
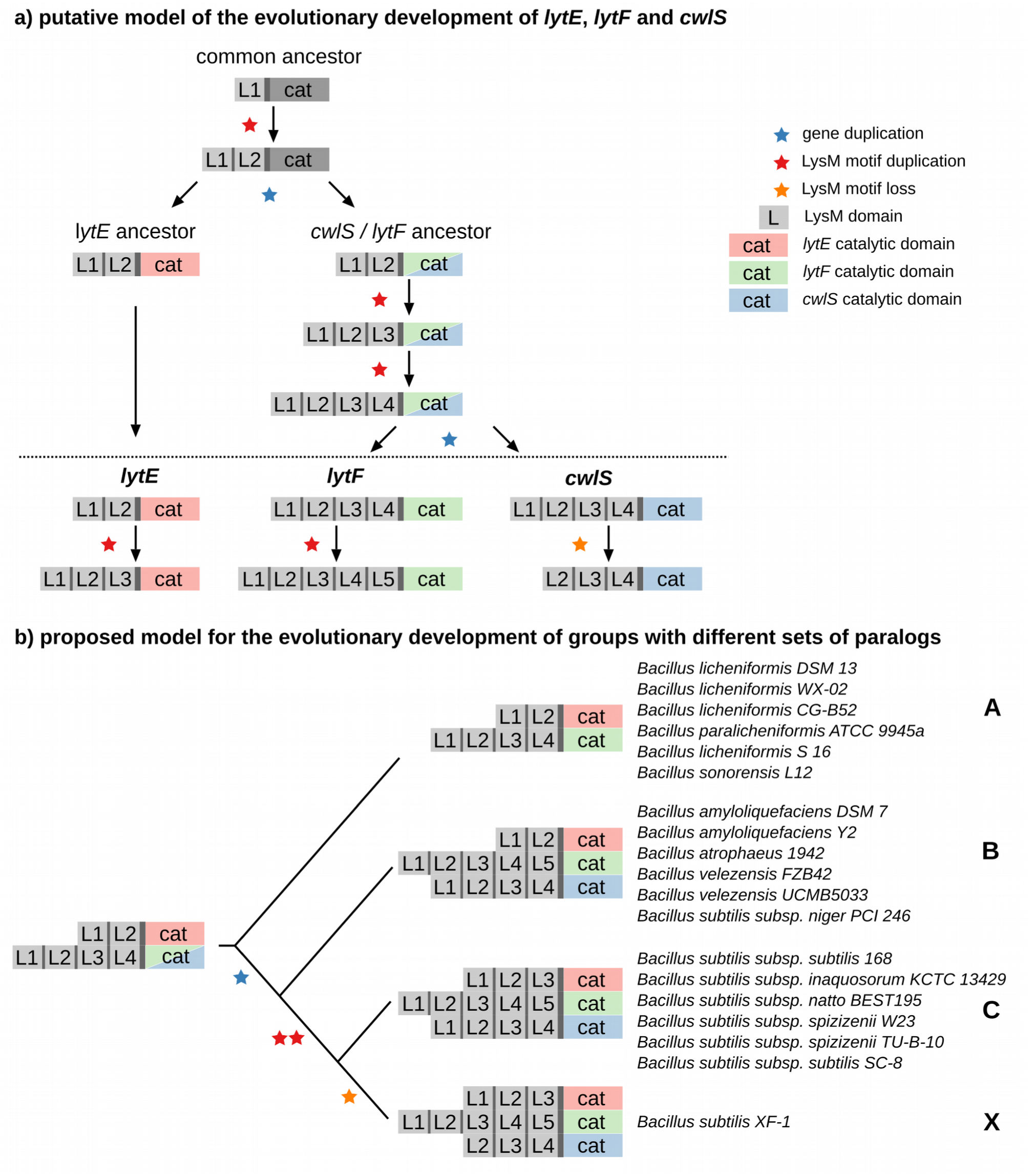
<u>6a: A putative model of the evolutionary development of the *cwlS*, *lytE* and *lytF* D,L- endopeptidases and their LysM motifs.</u> The proposed model assumes a stable inheritance of genes and LysM motifs. The common ancestor is a gene that encodes for a D,L-endopeptidase type protein with one LysM motif. Through a combination of gene duplication (blue star), LysM motif duplication (red star) as well as LysM motif loss events (orange star), the model arrives at the set of genes observed in this study, with LytE orthologs (having either two or three LysM motifs), CwlS orthologs (four or three LysM motifs) and LytF orthologs (with four or five LysM motifs). <u>6b: Proposed model for the evolutionary development of groups with different sets of paralogs.</u> The model assumes a stable inheritance of genes and LysM motifs and describes how through a combination of LysM motif (red star) and gene (blue star) duplication events as well as loss of LysM motifs (orange star) different bacterial species gain different sets of paralogs.

## Conclusion

Overall this study offers further insights into the D,L-endopeptidase requirement of *B. subtilis*. To our knowledge this is the first work that has demonstrated that the D,L- endopeptidase requirement of *B. subtilis* can be fulfilled by a D,L-endopeptidase other than LytE or CwlO. It remains unknown why LytF, but not CwlS is able to complement for CwlO, given their highly similar structure (also localization is a likely suspect).

This is also the first time that is has been investigated whether individual LysM domains are of equal importance for the functionality of an autolytic enzyme in vivo. It was shown that any single LysM domain is sufficient for functionality of the LytE enzyme, however robust growth requires the three LysM domains present in the wild type enzyme. Among these we demonstrated that the first LysM domain is superior over the subsequent domains, especially when located N-terminally, which is probably due to some interaction with the signaling peptide.

An investigation of bacterial LysM domains on a large scale led to some interesting observations, such as that expansion of LysM domains occurs frequently in the *Firmicutes* phylum and that LysM domains are unlikely to be transferred horizontally. Accordingly, a putative model was developed describing the relationship between the LytE, CwlS and LytF D,L-endopeptidases.

It is clear from our results that D,L-endopeptidases must have many features to be able to fulfill the requirement for viability. Among those requirements are the level of enzyme expressed, the stability of the protein when expressed and the localization of the protein within the cell envelope.

Further investigating these and other aspects of autolytic enzymes and cell wall metabolism will not only deepen our understanding of these processes, but also has the potential to be utilized for the development of new antibiotics and for the optimization of bacterial strains for enzyme and protein production. Furthermore an in-depth understanding of LysM domains and conditions that affect their biding affinity is likely to be of importance for the development of future binding assays and cell surface displays.

## Suplementary Material

**Supplementary Figure 1.**
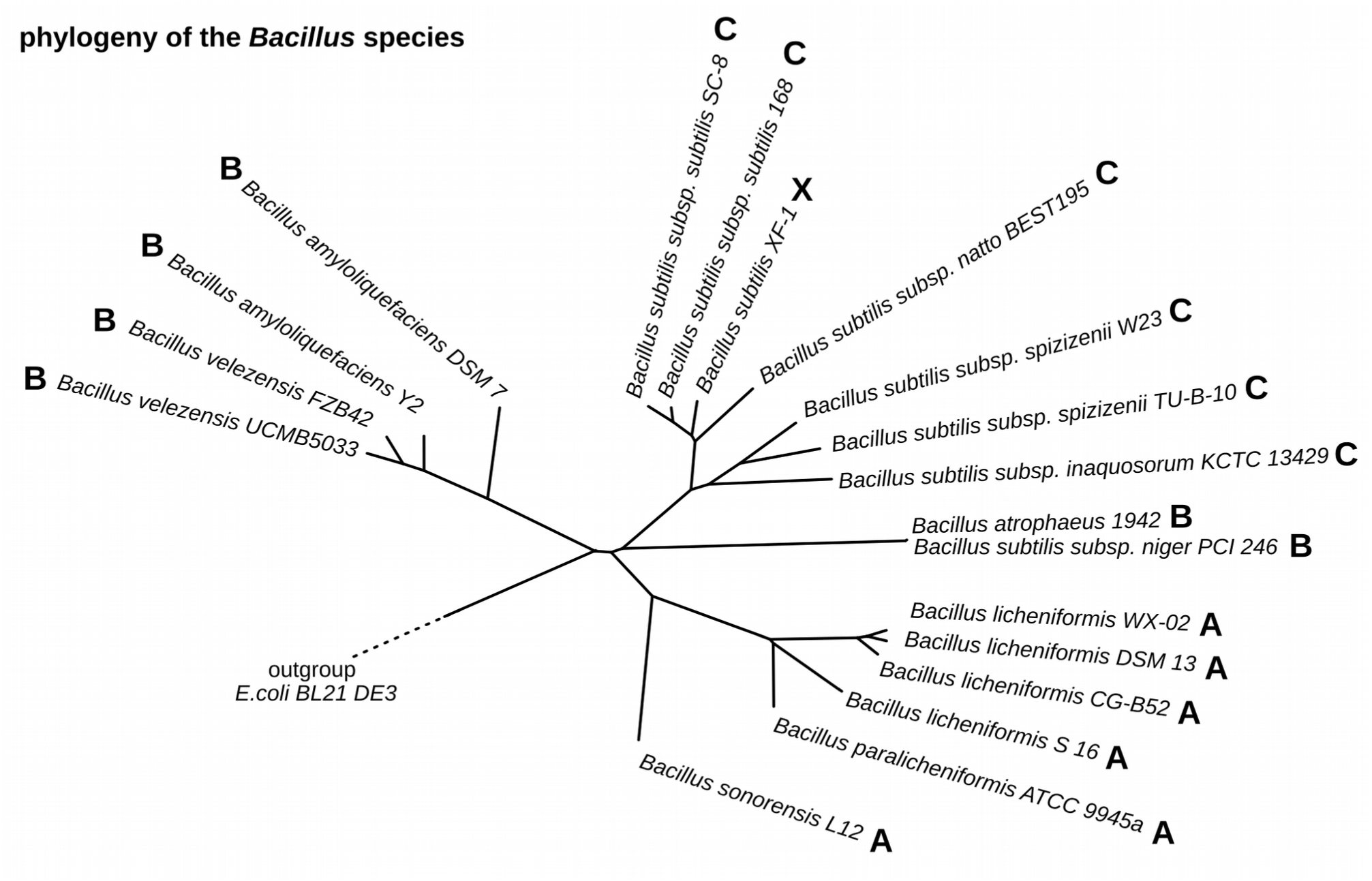
<u>Supplementary 1: Basic phylogeny of the *Bacillus* species.</u> Phylogeny of the investigated 19 Bacillus species was inferred through alignment-free genome comparison with feature frequency profiles (FFP). Genome sequences (excluding plasmid sequences) of all species were retrieved from GenBank. FPP was performed with length l = 24. A neighbour Joining (NJ) tree was constructed based on the Jensen-Shannon divergence matrice. Bootstrapping was performed to verify the robustness of the tree and boostrapping values of 100 out of 100 were achieved for all branches. Letters (A, B, C, X) refer to the different groups of paralogs defined in Figure 2b.

**Supplementary Figure 2:**
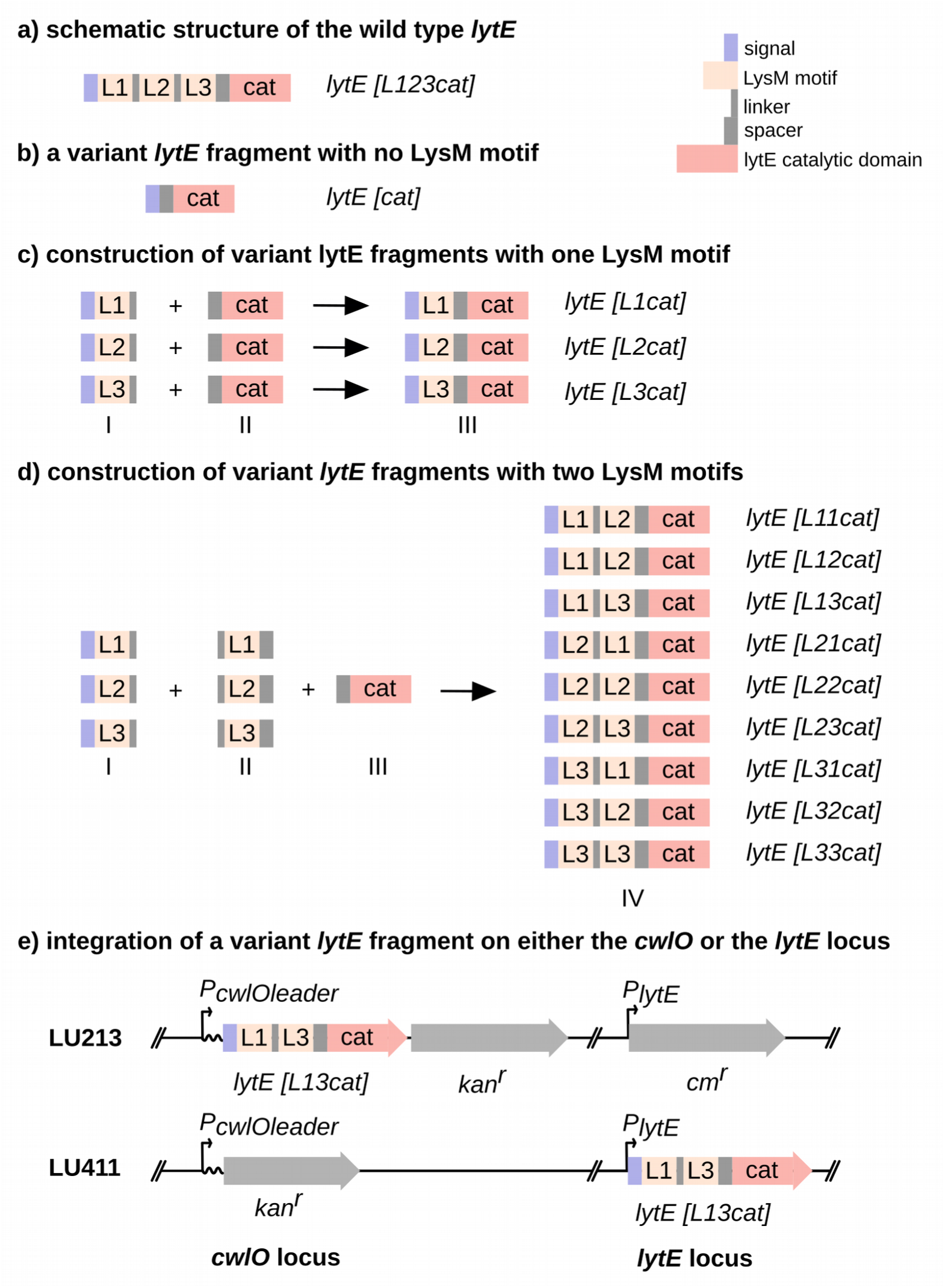
<u>Supplementary 2a: schematic structure of the wild type *lytE*</u> Schematic structure of the WT168 *lytE*, which features a signal sequence followed by three LysM motifs (L1, L2 and L3), separated by linker sequences and a catalytic domain (cat) which is separated from L3 by a spacer sequence. <u>Supplementary 2b: schematic structure of a variant *lytE* fragment with no LysM motifs</u> A linear fragment encoding a variant *lytE* without a LysM motif was generated by amplifying the catalytic domain with (forward) primers flanked by the signal sequence. <u>Supplementary 2c: construction of variant *lytE* fragments with one LysM motifs</u> Linear fragments encoding variant *lytE* with one LysM motif (III), are constructed by amplifying individual LysM motifs (I), with primers flanked by the signal and the spacer sequence, and joining them to the catalytic domain (II) by long-flanking homology PCR, thereby utilizing the spacer sequence for strand overlap. <u>Supplementary 2d: construction of variant *lytE* fragments with two LysM motifs</u> PCR fragments encoding variant *lytE* with two LysM motifs (IV), were constructed by amplifying individual LysM motifs with two set of primers (I and II) (one primer set is flanked by the signal and the linker sequence and the other by the linker and spacer sequence) and joining them alternating to the catalytic domain (III) by long-flanking homology PCR, thereby utilizing the linker and spacer sequence for strand overlap to generate all nine possible permutations. <u>Supplementary 2e: integration of a variant *lytE* fragment on either the *cwlO* or the *lytE* locus</u> Schematic representation of strains LU213 and LU411, in which a variant LytE open reading frame with 2 LysM motifs in the combination L1-L3 is integrated on the chromosome. In LU213 the variant LytE is expressed from the *cwlO* promoter with the *cwlO* leader sequence (replacing the endogenous *cwlO* gene) and in LU411 the variant *lytE* is expressed from from the *lytE* promoter (replacing the endogenous *lytE* gene). The endogenous *lytE* gene in LU213 is disrupted with a chloramphenicol resistance cassette, while the endogenous *cwlO* gene in LU411 is disrupted with kanamycin resistance cassette. Open reading frames for *lytE* and *cwlO* are not contiguous.

**Table S1:**
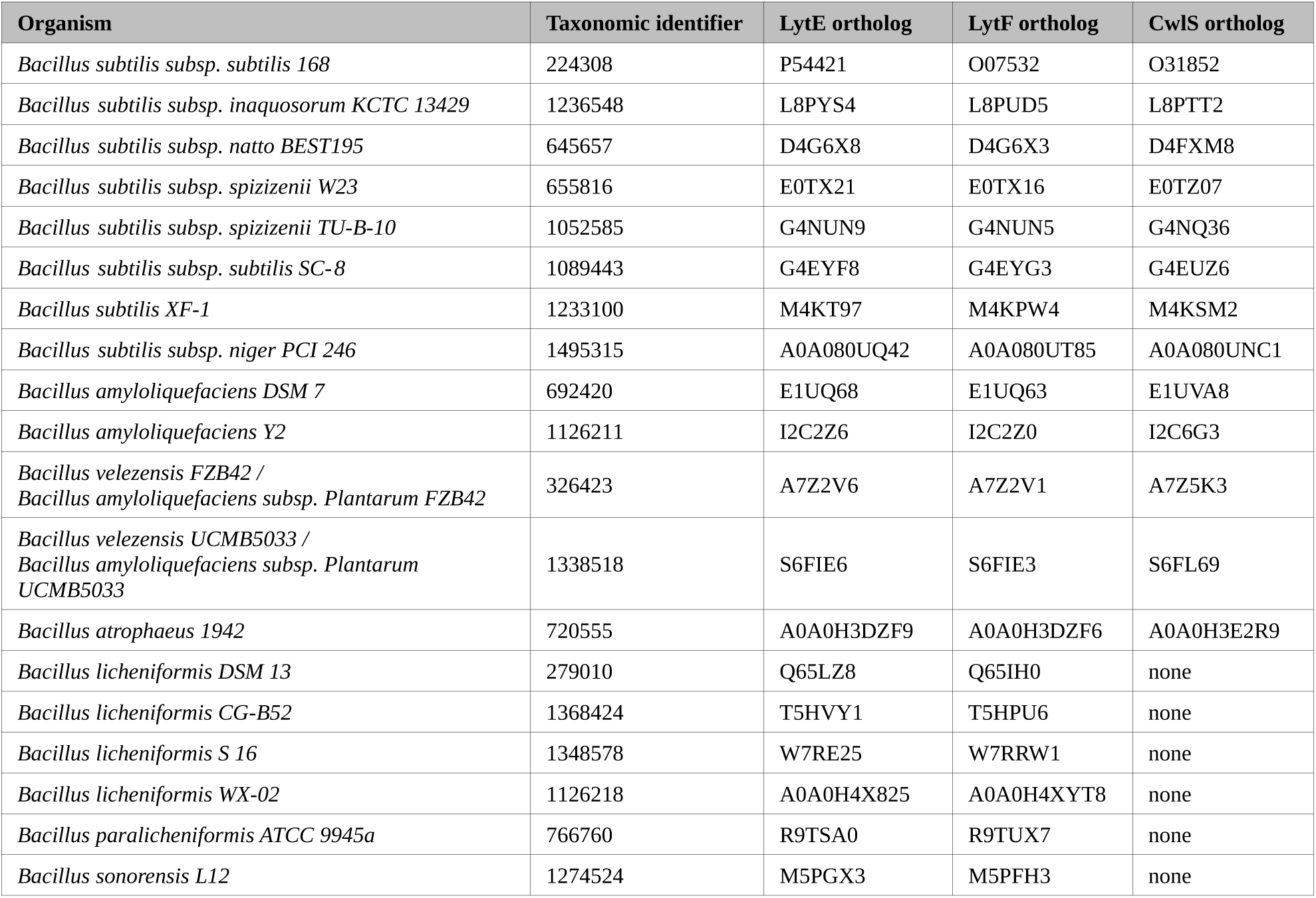
List of the identified orthologs to LytE, CwlS and LytF with their UniProt identifier and NCBI taxonomic identifier

**Table S2:**
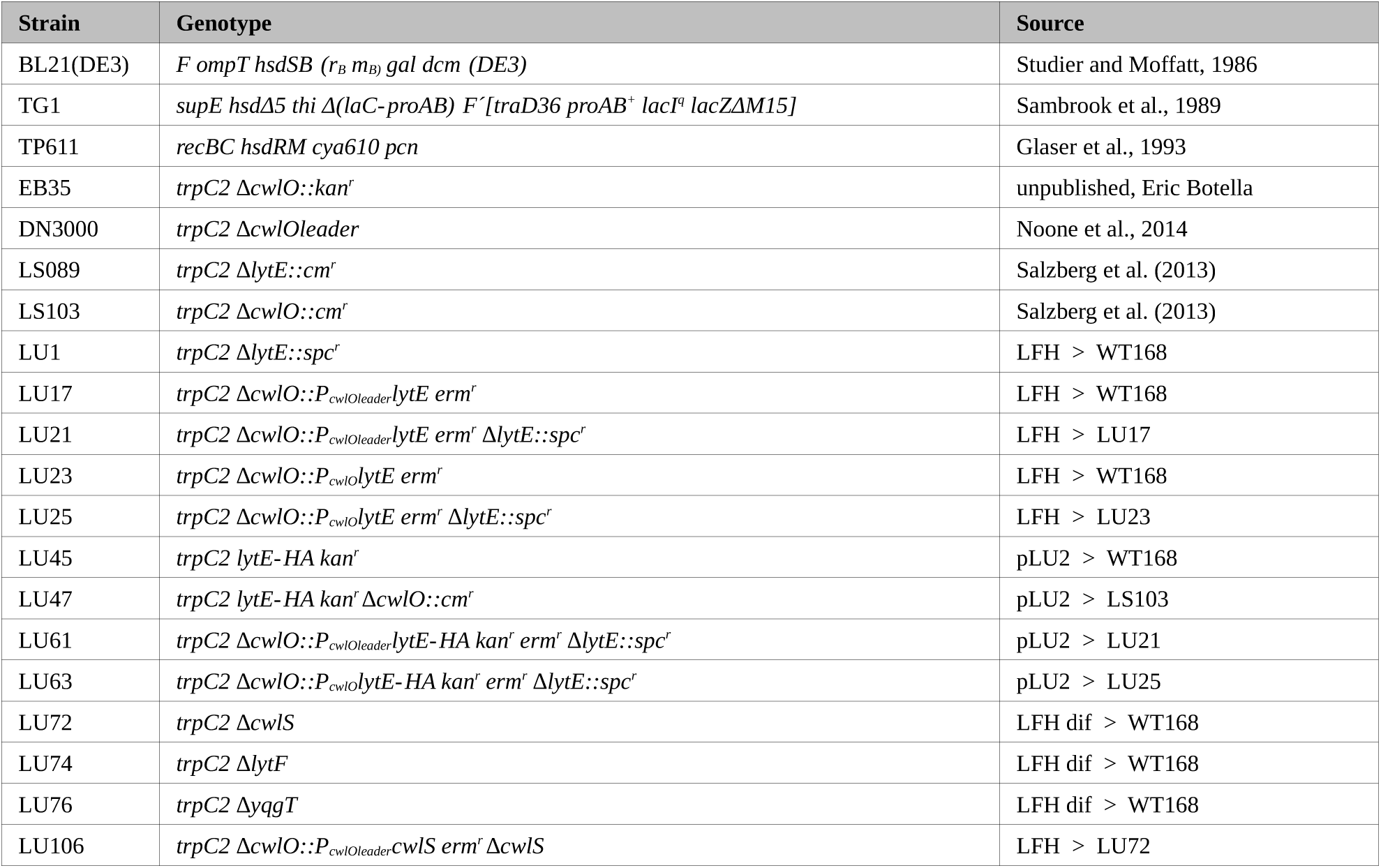
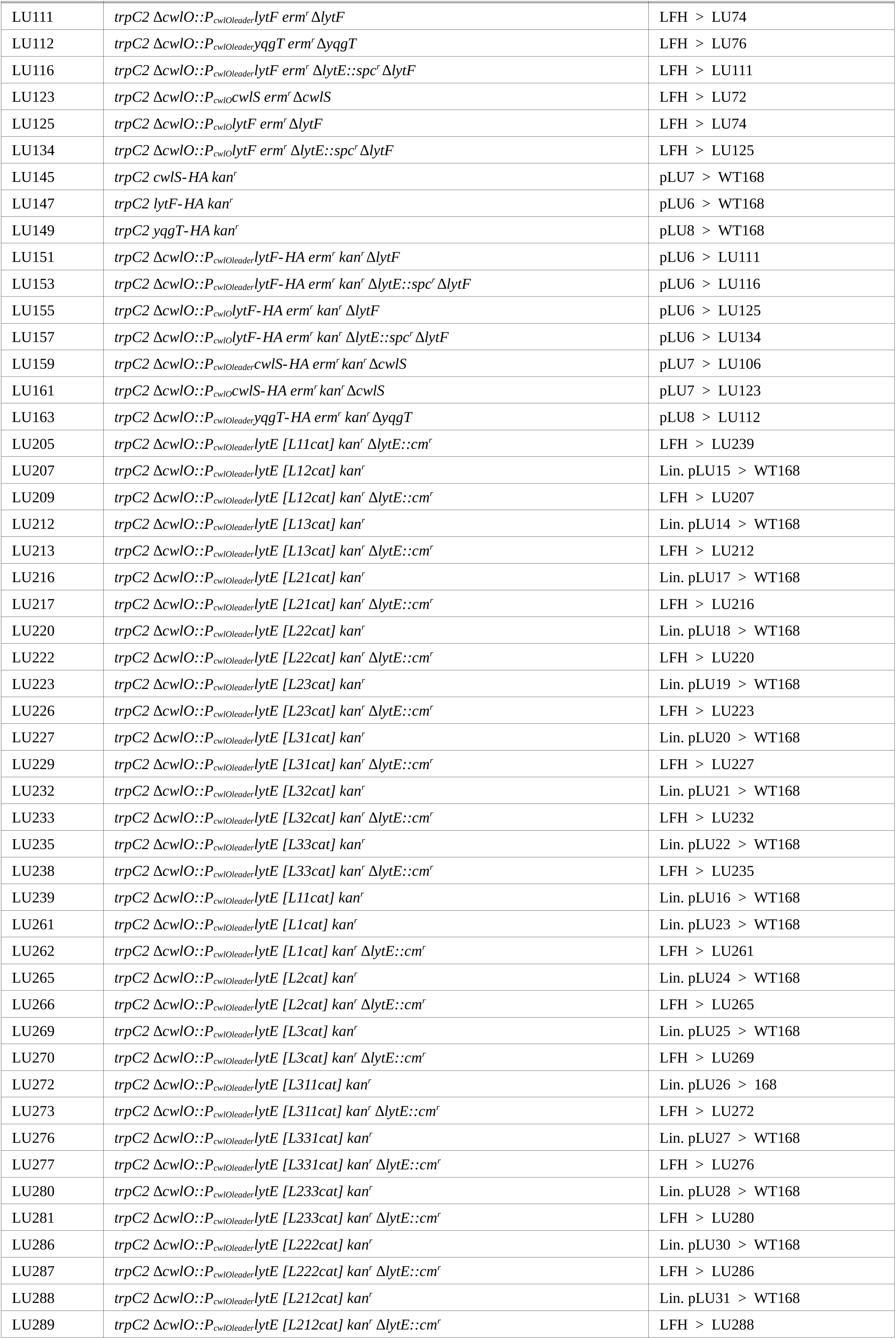
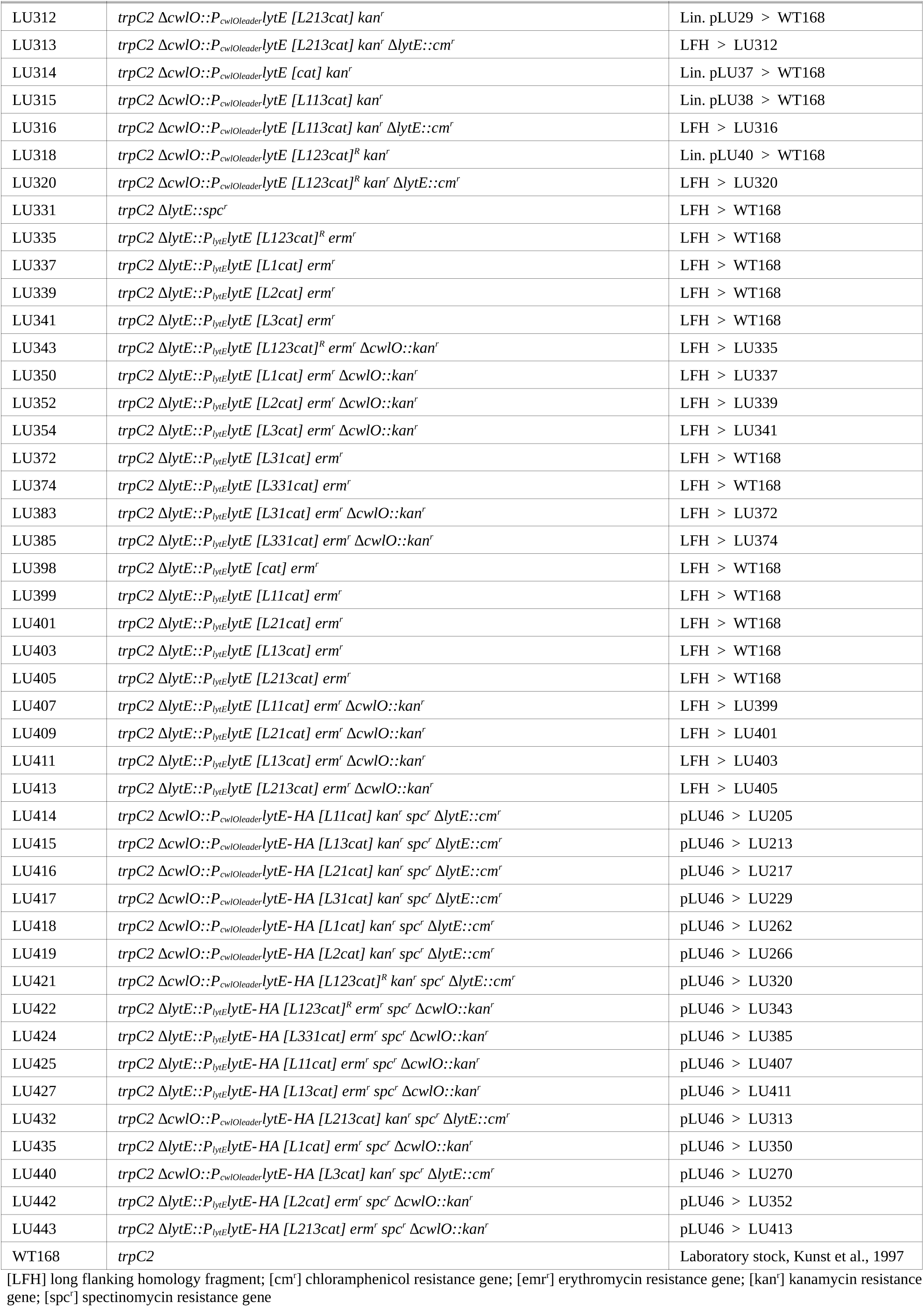
List of strains used in this study

| Strain | Genotype | Source |
| --- | --- | --- |
| BL21(DE3) | *F ompT hsdSB (r<sub>B</sub> m<sub>B</sub>) gal dcm (DE3)* | Studier and Moffatt, 1986 |
| TG1 | *supE hsdΔ5 thi Δ(lac-proAB) F'[traD36 proAB<sup>+</sup> lacI<sup>q</sup> lacZΔM15]* | Sambrook et al., 1989 |
| TP611 | *recBC hsdRM cya610 pcn* | Glaser et al., 1993 |
| EB35 | *trpC2 ΔcwlO::kan<sup>r</sup>* | unpublished, Eric Botella |
| DN3000 | *trpC2 ΔcwlOleader* | Noone et al., 2014 |
| LS089 | *trpC2 ΔlytE::cm<sup>r</sup>* | Salzberg et al. (2013) |
| LS103 | *trpC2 ΔcwlO::cm<sup>r</sup>* | Salzberg et al. (2013) |
| LU1 | *trpC2 ΔlytE::spc<sup>r</sup>* | LFH > WT168 |
| LU17 | *trpC2 ΔcwlO::P<sub>cwlOleader</sub>lytE erm<sup>r</sup>* | LFH > WT168 |
| LU21 | *trpC2 ΔcwlO::P<sub>cwlOleader</sub>lytE erm<sup>r</sup> ΔlytE::spc<sup>r</sup>* | LFH > LU17 |
| LU23 | *trpC2 ΔcwlO::P<sub>cwlO</sub>lytE erm<sup>r</sup>* | LFH > WT168 |
| LU25 | *trpC2 ΔcwlO::P<sub>cwlO</sub>lytE erm<sup>r</sup> ΔlytE::spc<sup>r</sup>* | LFH > LU23 |
| LU45 | *trpC2 lytE-HA kan<sup>r</sup>* | pLU2 > WT168 |
| LU47 | *trpC2 lytE-HA kan<sup>r</sup> ΔcwlO::cm<sup>r</sup>* | pLU2 > LS103 |
| LU61 | *trpC2 ΔcwlO::P<sub>cwlOleader</sub>lytE-HA kan<sup>r</sup> erm<sup>r</sup> ΔlytE::spc<sup>r</sup>* | pLU2 > LU21 |
| LU63 | *trpC2 ΔcwlO::P<sub>cwlO</sub>lytE-HA kan<sup>r</sup> erm<sup>r</sup> ΔlytE::spc<sup>r</sup>* | pLU2 > LU25 |
| LU72 | *trpC2 ΔcwlS* | LFH dif > WT168 |
| LU74 | *trpC2 ΔlytF* | LFH dif > WT168 |
| LU76 | *trpC2 ΔyqgT* | LFH dif > WT168 |
| LU106 | *trpC2 ΔcwlO::P<sub>cwlOleader</sub>CwlS erm<sup>r</sup> ΔcwlS* | LFH > LU72 |
| LU111 | *trpC2 ΔcwlO::P<sub>cwlO</sub>leader<sub>lytF</sub> erm<sup>r</sup> ΔlytF* | LFH > LU74 |
| LU112 | *trpC2 ΔcwlO::P<sub>cwlO</sub>leader<sub>yqgT</sub> erm<sup>r</sup> ΔyqgT* | LFH > LU76 |
| LU116 | *trpC2 ΔcwlO::P<sub>cwlO</sub>leader<sub>lytF</sub> erm<sup>r</sup> ΔlytE::spc<sup>r</sup> ΔlytF* | LFH > LU111 |
| LU123 | *trpC2 ΔcwlO::P<sub>cwlO</sub>cwlS erm<sup>r</sup> ΔcwlS* | LFH > LU72 |
| LU125 | *trpC2 ΔcwlO::P<sub>cwlO</sub>lytF erm<sup>r</sup> ΔlytF* | LFH > LU74 |
| LU134 | *trpC2 ΔcwlO::P<sub>cwlO</sub>lytF erm<sup>r</sup> ΔlytE::spc<sup>r</sup> ΔlytF* | LFH > LU125 |
| LU145 | *trpC2 cwlS-HA kan<sup>r</sup>* | pLU7 > WT168 |
| LU147 | *trpC2 lytF-HA kan<sup>r</sup>* | pLU6 > WT168 |
| LU149 | *trpC2 yqgT-HA kan<sup>r</sup>* | pLU8 > WT168 |
| LU151 | *trpC2 ΔcwlO::P<sub>cwlO</sub>leader<sub>lytF</sub>-HA erm<sup>r</sup> kan<sup>r</sup> ΔlytF* | pLU6 > LU111 |
| LU153 | *trpC2 ΔcwlO::P<sub>cwlO</sub>leader<sub>lytF</sub>-HA erm<sup>r</sup> kan<sup>r</sup> ΔlytE::spc<sup>r</sup> ΔlytF* | pLU6 > LU116 |
| LU155 | *trpC2 ΔcwlO::P<sub>cwlO</sub>lytF-HA erm<sup>r</sup> kan<sup>r</sup> ΔlytF* | pLU6 > LU125 |
| LU157 | *trpC2 ΔcwlO::P<sub>cwlO</sub>lytF-HA erm<sup>r</sup> kan<sup>r</sup> ΔlytE::spc<sup>r</sup> ΔlytF* | pLU6 > LU134 |
| LU159 | *trpC2 ΔcwlO::P<sub>cwlO</sub>leader<sub>cwlS</sub>-HA erm<sup>r</sup> kan<sup>r</sup> ΔcwlS* | pLU7 > LU106 |
| LU161 | *trpC2 ΔcwlO::P<sub>cwlO</sub>cwlS-HA erm<sup>r</sup> kan<sup>r</sup> ΔcwlS* | pLU7 > LU123 |
| LU163 | *trpC2 ΔcwlO::P<sub>cwlO</sub>leader<sub>yqgT</sub>-HA erm<sup>r</sup> kan<sup>r</sup> ΔyqgT* | pLU8 > LU112 |
| LU205 | *trpC2 ΔcwlO::P<sub>cwlO</sub>leader<sub>lytE</sub> [L11cat] kan<sup>r</sup> ΔlytE::cm<sup>r</sup>* | LFH > LU239 |
| LU207 | *trpC2 ΔcwlO::P<sub>cwlO</sub>leader<sub>lytE</sub> [L12cat] kan<sup>r</sup>* | Lin. pLU15 > WT168 |
| LU209 | *trpC2 ΔcwlO::P<sub>cwlO</sub>leader<sub>lytE</sub> [L12cat] kan<sup>r</sup> ΔlytE::cm<sup>r</sup>* | LFH > LU207 |
| LU212 | *trpC2 ΔcwlO::P<sub>cwlO</sub>leader<sub>lytE</sub> [L13cat] kan<sup>r</sup>* | Lin. pLU14 > WT168 |
| LU213 | *trpC2 ΔcwlO::P<sub>cwlO</sub>leader<sub>lytE</sub> [L13cat] kan<sup>r</sup> ΔlytE::cm<sup>r</sup>* | LFH > LU212 |
| LU216 | *trpC2 ΔcwlO::P<sub>cwlO</sub>leader<sub>lytE</sub> [L21cat] kan<sup>r</sup>* | Lin. pLU17 > WT168 |
| LU217 | *trpC2 ΔcwlO::P<sub>cwlO</sub>leader<sub>lytE</sub> [L21cat] kan<sup>r</sup> ΔlytE::cm<sup>r</sup>* | LFH > LU216 |
| LU220 | *trpC2 ΔcwlO::P<sub>cwlO</sub>leader<sub>lytE</sub> [L22cat] kan<sup>r</sup>* | Lin. pLU18 > WT168 |
| LU222 | *trpC2 ΔcwlO::P<sub>cwlO</sub>leader<sub>lytE</sub> [L22cat] kan<sup>r</sup> ΔlytE::cm<sup>r</sup>* | LFH > LU220 |
| LU223 | *trpC2 ΔcwlO::P<sub>cwlO</sub>leader<sub>lytE</sub> [L23cat] kan<sup>r</sup>* | Lin. pLU19 > WT168 |
| LU226 | *trpC2 ΔcwlO::P<sub>cwlO</sub>leader<sub>lytE</sub> [L23cat] kan<sup>r</sup> ΔlytE::cm<sup>r</sup>* | LFH > LU223 |
| LU227 | *trpC2 ΔcwlO::P<sub>cwlO</sub>leader<sub>lytE</sub> [L31cat] kan<sup>r</sup>* | Lin. pLU20 > WT168 |
| LU229 | *trpC2 ΔcwlO::P<sub>cwlO</sub>leader<sub>lytE</sub> [L31cat] kan<sup>r</sup> ΔlytE::cm<sup>r</sup>* | LFH > LU227 |
| LU232 | *trpC2 ΔcwlO::P<sub>cwlO</sub>leader<sub>lytE</sub> [L32cat] kan<sup>r</sup>* | Lin. pLU21 > WT168 |
| LU233 | *trpC2 ΔcwlO::P<sub>cwlO</sub>leader<sub>lytE</sub> [L32cat] kan<sup>r</sup> ΔlytE::cm<sup>r</sup>* | LFH > LU232 |
| LU235 | *trpC2 ΔcwlO::P<sub>cwlO</sub>leader<sub>lytE</sub> [L33cat] kan<sup>r</sup>* | Lin. pLU22 > WT168 |
| LU238 | *trpC2 ΔcwlO::P<sub>cwlO</sub>leader<sub>lytE</sub> [L33cat] kan<sup>r</sup> ΔlytE::cm<sup>r</sup>* | LFH > LU235 |
| LU239 | *trpC2 ΔcwlO::P<sub>cwlO</sub>leader<sub>lytE</sub> [L11cat] kan<sup>r</sup>* | Lin. pLU16 > WT168 |
| LU261 | *trpC2 ΔcwlO::P<sub>cwlO</sub>leader<sub>lytE</sub> [L1cat] kan<sup>r</sup>* | Lin. pLU23 > WT168 |
| LU262 | *trpC2 ΔcwlO::P<sub>cwlO</sub>leader<sub>lytE</sub> [L1cat] kan<sup>r</sup> ΔlytE::cm<sup>r</sup>* | LFH > LU261 |
| LU265 | *trpC2 ΔcwlO::P<sub>cwlO</sub>leader<sub>lytE</sub> [L2cat] kan<sup>r</sup>* | Lin. pLU24 > WT168 |
| LU266 | *trpC2 ΔcwlO::P<sub>cwlO</sub>leader<sub>lytE</sub> [L2cat] kan<sup>r</sup> ΔlytE::cm<sup>r</sup>* | LFH > LU265 |
| LU269 | *trpC2 ΔcwlO::P<sub>cwlO</sub>leader<sub>lytE</sub> [L3cat] kan<sup>r</sup>* | Lin. pLU25 > WT168 |
| LU270 | *trpC2 ΔcwlO::P<sub>cwlO</sub>leader<sub>lytE</sub> [L3cat] kan<sup>r</sup> ΔlytE::cm<sup>r</sup>* | LFH > LU269 |
| LU272 | *trpC2 ΔcwlO::P<sub>cwlO</sub>leader<sub>lytE</sub> [L311cat] kan<sup>r</sup>* | Lin. pLU26 > 168 |
| LU273 | *trpC2 ΔcwlO::P<sub>cwlO</sub>leader<sub>lytE</sub> [L311cat] kan<sup>r</sup> ΔlytE::cm<sup>r</sup>* | LFH > LU272 |
| LU276 | *trpC2 ΔcwlO::P<sub>cwlO</sub>leader<sub>lytE</sub> [L331cat] kan<sup>r</sup>* | Lin. pLU27 > WT168 |
| LU277 | *trpC2 ΔcwlO::P<sub>cwlO</sub>leader<sub>lytE</sub> [L331cat] kan<sup>r</sup> ΔlytE::cm<sup>r</sup>* | LFH > LU276 |
| LU280 | *trpC2 ΔcwlO::P<sub>cwlO</sub>leader<sub>lytE</sub> [L233cat] kan<sup>r</sup>* | Lin. pLU28 > WT168 |
| LU281 | *trpC2 ΔcwlO::P<sub>cwlO</sub>leader<sub>lytE</sub> [L233cat] kan<sup>r</sup> ΔlytE::cm<sup>r</sup>* | LFH > LU280 |
| LU286 | *trpC2 ΔcwlO::P<sub>cwlO</sub>leader<sub>lytE</sub> [L222cat] kan<sup>r</sup>* | Lin. pLU30 > WT168 |
| LU287 | *trpC2 ΔcwlO::P<sub>cwlO</sub>leader<sub>lytE</sub> [L222cat] kan<sup>r</sup> ΔlytE::cm<sup>r</sup>* | LFH > LU286 |
| LU288 | *trpC2 ΔcwlO::P<sub>cwlO</sub>leader<sub>lytE</sub> [L212cat] kan<sup>r</sup>* | Lin. pLU31 > WT168 |
| LU289 | *trpC2 ΔcwlO::P<sub>cwlO</sub>leader<sub>lytE</sub> [L212cat] kan<sup>r</sup> ΔlytE::cm<sup>r</sup>* | LFH > LU288 |
| LU312 | *trpC2 ΔcwlO::P<sub>cwlO</sub>leader<sub>lytE</sub> [L213cat] kan<sup>r</sup>* | Lin. pLU29 > WT168 |
| LU313 | *trpC2 ΔcwlO::P<sub>cwlO</sub>leader<sub>lytE</sub> [L213cat] kan<sup>r</sup> ΔlytE::cm<sup>r</sup>* | LFH > LU312 |
| LU314 | *trpC2 ΔcwlO::P<sub>cwlO</sub>leader<sub>lytE</sub> [cat] kan<sup>r</sup>* | Lin. pLU37 > WT168 |
| LU315 | *trpC2 ΔcwlO::P<sub>cwlO</sub>leader<sub>lytE</sub> [L113cat] kan<sup>r</sup>* | Lin. pLU38 > WT168 |
| LU316 | *trpC2 ΔcwlO::P<sub>cwlO</sub>leader<sub>lytE</sub> [L113cat] kan<sup>r</sup> ΔlytE::cm<sup>r</sup>* | LFH > LU316 |
| LU318 | *trpC2 ΔcwlO::P<sub>cwlO</sub>leader<sub>lytE</sub> [L123cat]<sup>R</sup> kan<sup>r</sup>* | Lin. pLU40 > WT168 |
| LU320 | *trpC2 ΔcwlO::P<sub>cwlO</sub>leader<sub>lytE</sub> [L123cat]<sup>R</sup> kan<sup>r</sup> ΔlytE::cm<sup>r</sup>* | LFH > LU320 |
| LU331 | *trpC2 ΔlytE::spc<sup>r</sup>* | LFH > WT168 |
| LU335 | *trpC2 ΔlytE::P<sub>lytE</sub><sub>lytE</sub> [L123cat]<sup>R</sup> erm<sup>r</sup>* | LFH > WT168 |
| LU337 | *trpC2 ΔlytE::P<sub>lytE</sub><sub>lytE</sub> [L1cat] erm<sup>r</sup>* | LFH > WT168 |
| LU339 | *trpC2 ΔlytE::P<sub>lytE</sub><sub>lytE</sub> [L2cat] erm<sup>r</sup>* | LFH > WT168 |
| LU341 | *trpC2 ΔlytE::P<sub>lytE</sub><sub>lytE</sub> [L3cat] erm<sup>r</sup>* | LFH > WT168 |
| LU343 | *trpC2 ΔlytE::P<sub>lytE</sub><sub>lytE</sub> [L123cat]<sup>R</sup> erm<sup>r</sup> ΔcwlO::kan<sup>r</sup>* | LFH > LU335 |
| LU350 | *trpC2 ΔlytE::P<sub>lytE</sub><sub>lytE</sub> [L1cat] erm<sup>r</sup> ΔcwlO::kan<sup>r</sup>* | LFH > LU337 |
| LU352 | *trpC2 ΔlytE::P<sub>lytE</sub><sub>lytE</sub> [L2cat] erm<sup>r</sup> ΔcwlO::kan<sup>r</sup>* | LFH > LU339 |
| LU354 | *trpC2 ΔlytE::P<sub>lytE</sub><sub>lytE</sub> [L3cat] erm<sup>r</sup> ΔcwlO::kan<sup>r</sup>* | LFH > LU341 |
| LU372 | *trpC2 ΔlytE::P<sub>lytE</sub><sub>lytE</sub> [L31cat] erm<sup>r</sup>* | LFH > WT168 |
| LU374 | *trpC2 ΔlytE::P<sub>lytE</sub><sub>lytE</sub> [L331cat] erm<sup>r</sup>* | LFH > WT168 |
| LU383 | *trpC2 ΔlytE::P<sub>lytE</sub><sub>lytE</sub> [L31cat] erm<sup>r</sup> ΔcwlO::kan<sup>r</sup>* | LFH > LU372 |
| LU385 | *trpC2 ΔlytE::P<sub>lytE</sub><sub>lytE</sub> [L331cat] erm<sup>r</sup> ΔcwlO::kan<sup>r</sup>* | LFH > LU374 |
| LU398 | *trpC2 ΔlytE::P<sub>lytE</sub><sub>lytE</sub> [cat] erm<sup>r</sup>* | LFH > WT168 |
| LU399 | *trpC2 ΔlytE::P<sub>lytE</sub><sub>lytE</sub> [L11cat] erm<sup>r</sup>* | LFH > WT168 |
| LU401 | *trpC2 ΔlytE::P<sub>lytE</sub><sub>lytE</sub> [L21cat] erm<sup>r</sup>* | LFH > WT168 |
| LU403 | *trpC2 ΔlytE::P<sub>lytE</sub><sub>lytE</sub> [L13cat] erm<sup>r</sup>* | LFH > WT168 |
| LU405 | *trpC2 ΔlytE::P<sub>lytE</sub><sub>lytE</sub> [L213cat] erm<sup>r</sup>* | LFH > WT168 |
| LU407 | *trpC2 ΔlytE::P<sub>lytE</sub><sub>lytE</sub> [L11cat] erm<sup>r</sup> ΔcwlO::kan<sup>r</sup>* | LFH > LU399 |
| LU409 | *trpC2 ΔlytE::P<sub>lytE</sub><sub>lytE</sub> [L21cat] erm<sup>r</sup> ΔcwlO::kan<sup>r</sup>* | LFH > LU401 |
| LU411 | *trpC2 ΔlytE::P<sub>lytE</sub><sub>lytE</sub> [L13cat] erm<sup>r</sup> ΔcwlO::kan<sup>r</sup>* | LFH > LU403 |
| LU413 | *trpC2 ΔlytE::P<sub>lytE</sub><sub>lytE</sub> [L213cat] erm<sup>r</sup> ΔcwlO::kan<sup>r</sup>* | LFH > LU405 |
| LU414 | *trpC2 ΔcwlO::P<sub>cwlO</sub>leader<sub>lytE</sub>-HA [L11cat] kan<sup>r</sup> spc<sup>r</sup> ΔlytE::cm<sup>r</sup>* | pLU46 > LU205 |
| LU415 | *trpC2 ΔcwlO::P<sub>cwlO</sub>leader<sub>lytE</sub>-HA [L13cat] kan<sup>r</sup> spc<sup>r</sup> ΔlytE::cm<sup>r</sup>* | pLU46 > LU213 |
| LU416 | *trpC2 ΔcwlO::P<sub>cwlO</sub>leader<sub>lytE</sub>-HA [L21cat] kan<sup>r</sup> spc<sup>r</sup> ΔlytE::cm<sup>r</sup>* | pLU46 > LU217 |
| LU417 | *trpC2 ΔcwlO::P<sub>cwlO</sub>leader<sub>lytE</sub>-HA [L31cat] kan<sup>r</sup> spc<sup>r</sup> ΔlytE::cm<sup>r</sup>* | pLU46 > LU229 |
| LU418 | *trpC2 ΔcwlO::P<sub>cwlO</sub>leader<sub>lytE</sub>-HA [L1cat] kan<sup>r</sup> spc<sup>r</sup> ΔlytE::cm<sup>r</sup>* | pLU46 > LU262 |
| LU419 | *trpC2 ΔcwlO::P<sub>cwlO</sub>leader<sub>lytE</sub>-HA [L2cat] kan<sup>r</sup> spc<sup>r</sup> ΔlytE::cm<sup>r</sup>* | pLU46 > LU266 |
| LU421 | *trpC2 ΔcwlO::P<sub>cwlO</sub>leader<sub>lytE</sub>-HA [L123cat]<sup>R</sup> kan<sup>r</sup> spc<sup>r</sup> ΔlytE::cm<sup>r</sup>* | pLU46 > LU320 |
| LU422 | *trpC2 ΔlytE::P<sub>lytE</sub><sub>lytE</sub>-HA [L123cat]<sup>R</sup> erm<sup>r</sup> spc<sup>r</sup> ΔcwlO::kan<sup>r</sup>* | pLU46 > LU343 |
| LU424 | *trpC2 ΔlytE::P<sub>lytE</sub><sub>lytE</sub>-HA [L331cat] erm<sup>r</sup> spc<sup>r</sup> ΔcwlO::kan<sup>r</sup>* | pLU46 > LU385 |
| LU425 | *trpC2 ΔlytE::P<sub>lytE</sub><sub>lytE</sub>-HA [L11cat] erm<sup>r</sup> spc<sup>r</sup> ΔcwlO::kan<sup>r</sup>* | pLU46 > LU407 |
| LU427 | *trpC2 ΔlytE::P<sub>lytE</sub><sub>lytE</sub>-HA [L13cat] erm<sup>r</sup> spc<sup>r</sup> ΔcwlO::kan<sup>r</sup>* | pLU46 > LU411 |
| LU432 | *trpC2 ΔcwlO::P<sub>cwlO</sub>leader<sub>lytE</sub>-HA [L213cat] kan<sup>r</sup> spc<sup>r</sup> ΔlytE::cm<sup>r</sup>* | pLU46 > LU313 |
| LU435 | *trpC2 ΔlytE::P<sub>lytE</sub><sub>lytE</sub>-HA [L1cat] erm<sup>r</sup> spc<sup>r</sup> ΔcwlO::kan<sup>r</sup>* | pLU46 > LU350 |
| LU440 | *trpC2 ΔcwlO::P<sub>cwlO</sub>leader<sub>lytE</sub>-HA [L3cat] kan<sup>r</sup> spc<sup>r</sup> ΔlytE::cm<sup>r</sup>* | pLU46 > LU270 |
| LU442 | *trpC2 ΔlytE::P<sub>lytE</sub><sub>lytE</sub>-HA [L2cat] erm<sup>r</sup> spc<sup>r</sup> ΔcwlO::kan<sup>r</sup>* | pLU46 > LU352 |
| LU443 | *trpC2 ΔlytE::P<sub>lytE</sub><sub>lytE</sub>-HA [L213cat] erm<sup>r</sup> spc<sup>r</sup> ΔcwlO::kan<sup>r</sup>* | pLU46 > LU413 |
| WT168 | *trpC2* | Laboratory stock, Kunst et al., 1997 |

[LFH] long flanking homology fragment; [cm<sup>r</sup>] chloramphenicol resistance gene; [erm<sup>r</sup>] erythromycin resistance gene; [kan<sup>r</sup>] kanamycin resistance gene; [spc<sup>r</sup>] spectinomycin resistance gene

**Table S3:**
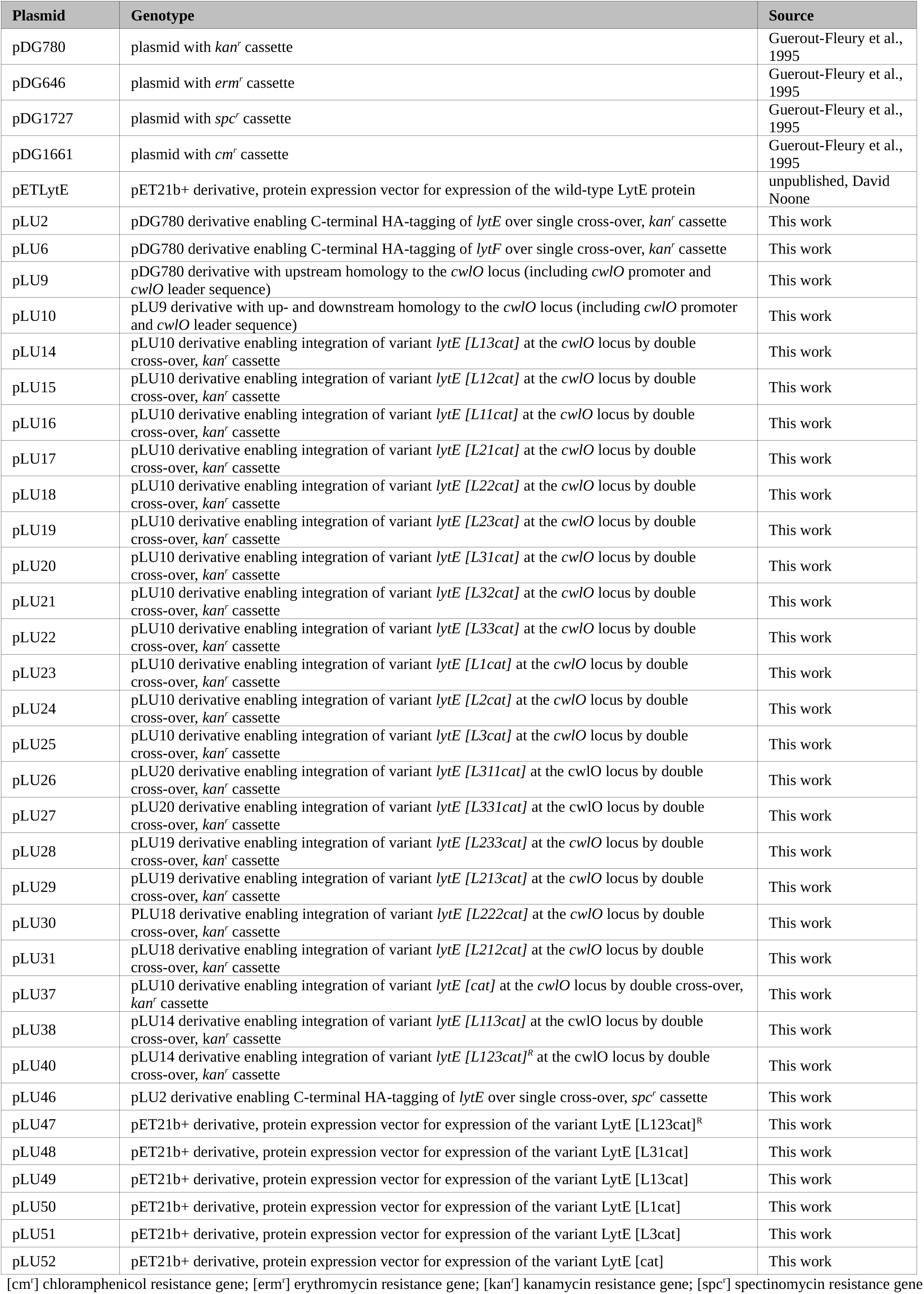
List of plasmids used in this study

**Table S4:**
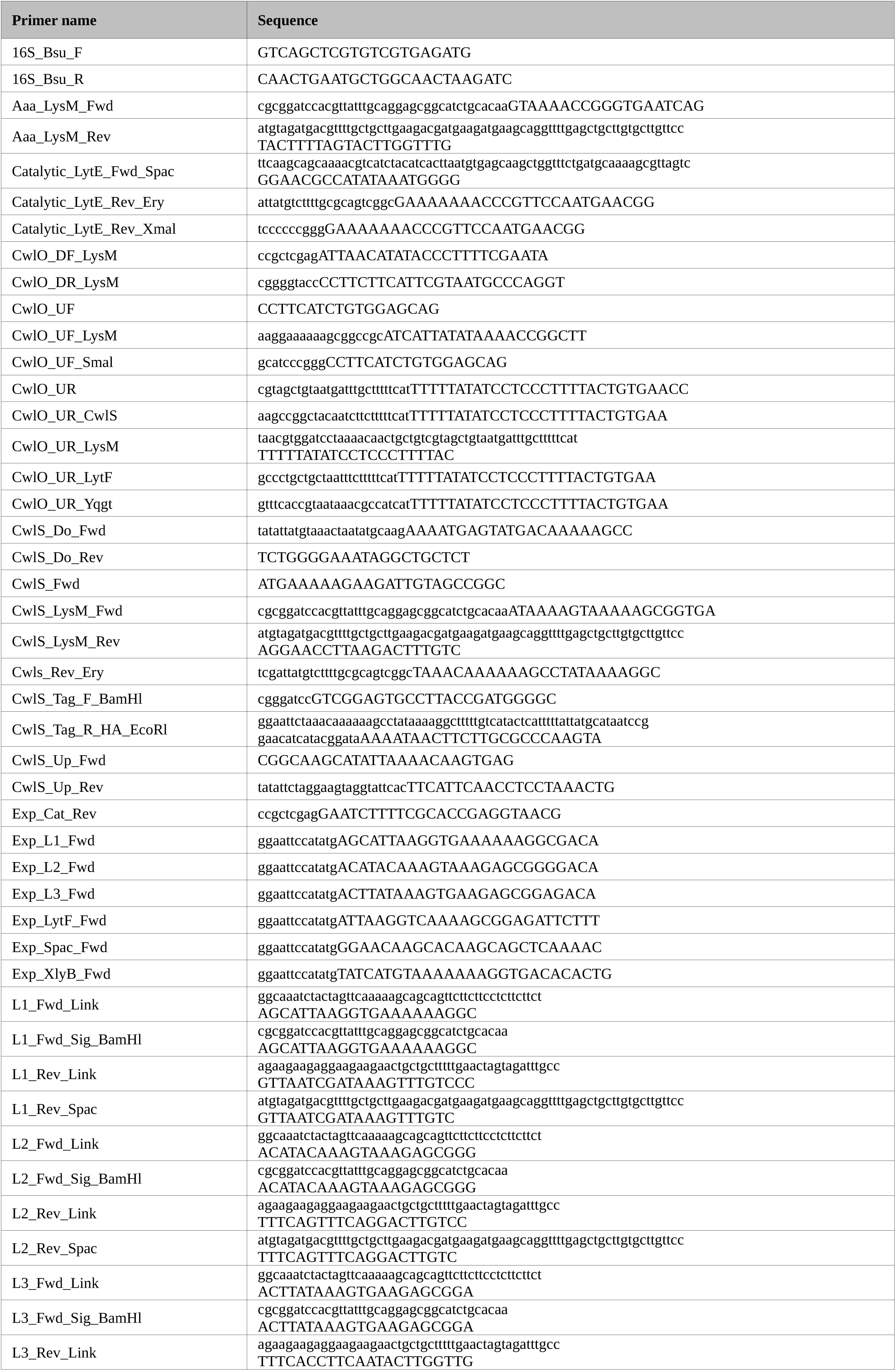
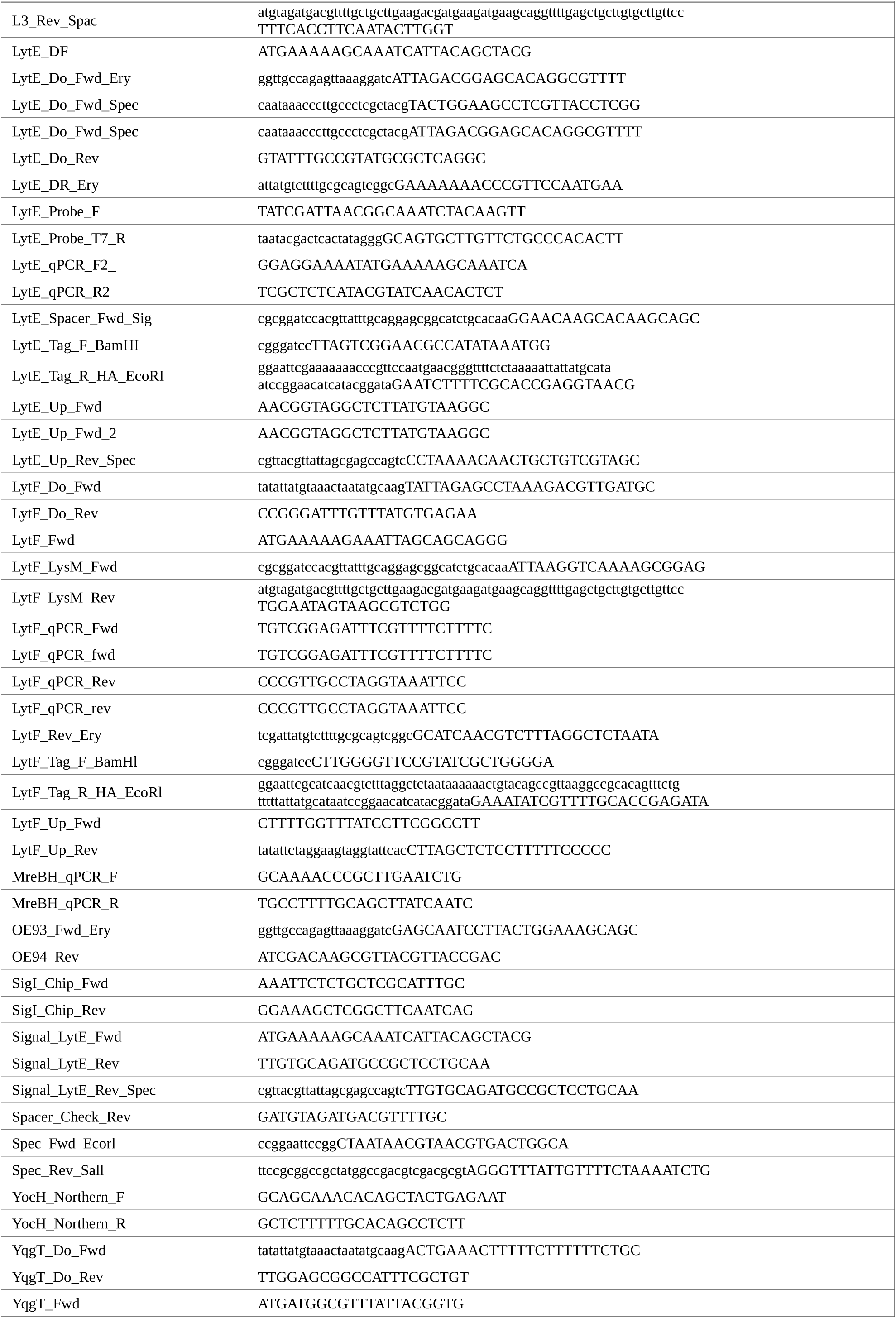
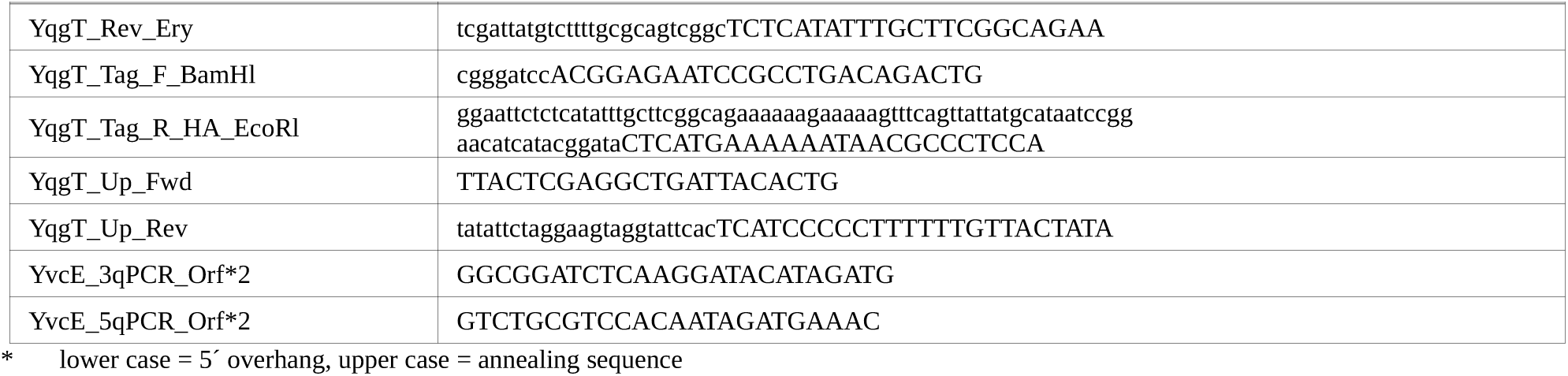
List of oligonucleotids used in this study

